# Evolutionary Trade-off Between Stomatal Defense and Gas Exchange in Brassicaceae

**DOI:** 10.1101/2025.03.31.646295

**Authors:** Wenshang Kang, Masahito Nakano, Kaori Fukumoto, Rikako Hirata, Pei Zhai, Yulin Du, Kunqi Hong, Jörg Ziegler, Yayoi Tsuda, Dieter Becker, Eiji Nambara, Ryohei Thomas Nakano, Shunsuke Miyashima, Akira Mine, Xiaowei Han, Kenichi Tsuda

**Author notes:** These authors contributed equally (Wenshang Kang, Masahito Nakano, Kaori Fukumoto, Rikako Hirata).

## Abstract

Stomatal opening is crucial for gas exchange, but it unavoidably offers invasion by pathogens. In response, plants close stomata to prevent pathogen entry, while the bacterial pathogen *Pseudomonas syringae* pv. tomato DC3000 (*Pto*) produces coronatine (COR), a jasmonate (JA) mimic, to counteract this plant response. Here, we demonstrate that by COR, *Pto* exploits *CYP707A1* activation in *Arabidopsis thaliana*, encoding an enzyme that degrades abscisic acid, essential for stomatal closure. Notably, COR-induction of *CYP707A1* is absent in other Brassicaceae species, such as *Capsella rubella* and *Eutrema salsugineum*, rendering them resistant to *Pto* invasion. In contrast, in *A. thaliana*, *CYP707A1* enables rapid stomatal opening at dawn in a JA-dependent manner, enhancing gas exchange and chlorophyll content, unlike *C. rubella* and *E. salsugineum*. Promoter-swap experiments confirm that the regulatory region of *CYP707A1* underlies these evolutionary diversifications. Together, our study presents a mechanism underlying an evolutionary trade-off between stomatal defense and gas exchange.

## INTRODUCTION

Stomata are pores on the plant leaf epidermis, formed by pairs of guard cells that sense and respond to internal and external cues.^1^ Guard cells regulate their turgor pressure to dynamically control the stomatal aperture, a key feature for plant survival and growth.^2,3^ One primary function of stomata is to mediate gas exchange between plants and the environment, by facilitating carbon dioxide uptake for photosynthesis and releasing water vapor through transpiration, which aids nutrient uptake from roots.^2,3^ The opening of stomata is vital for plant growth but poses a vulnerability, as it allows microbial pathogens to enter the plant through these pores, highlighting a significant trade-off between growth and defense.^4^ The balance between growth and defense is a critical adaptive challenge, particularly in wild plant species.^5^

The growth-defense trade-off has profound implications for agriculture. For example, the overexpression of plant resistance genes often enhances pathogen resistance at the expense of growth.^6^ This phenomenon can be attributed to either resource allocation constraints or genetically programmed regulatory mechanisms.^6^ Trade-offs are also central to evolutionary biology, as organisms optimize specific trade-off states that confer fitness in their environments and are regulated at various mechanisms, such as transcriptional regulation. For instance, in *Drosophila*, a single nucleotide polymorphism in the *cis*-regulatory region of *eyeless*, located within the binding site of its repressor, modifies temporal gene regulation and eye size, thereby mediating the trade-off between visual and olfactory organ development among *Drosophila* species.^7^ Suboptimal enhancer DNA sequences for transcription factor binding suggest an evolutionary trade-off between activity and specificity at multiple levels in eukaryotic transcriptional control.^8^ Furthermore, experimental evolution has shown that shifts in trade-offs frequently occur, driven by gene loss or regulatory changes.^9,10^ However, the evolutionary mechanisms underlying these shifts in growth-defense trade-offs in plants remain poorly understood.

Plants have evolved stomatal immunity as a defense mechanism against stomata-invading pathogens, closing their stomata upon recognizing Pathogen/Microbe-Associated Molecular Patterns (PAMPs/MAMPs), such as flg22, a bacterial flagellin component.^11^ Stomatal closure is mediated by pathways involving RBOH-dependent reactive oxygen species (ROS) production, calcium influx, MAP kinases, and the defense hormone salicylic acid (SA).^3,4^ Under drought conditions, stomatal closure is also regulated by the abscisic acid (ABA) signaling pathway, with overlapping components such as ABA biosynthesis, OST1 kinase, and the SLAC1 anion channel being critical for both ABA- and MAMP-induced closure.^4,12^

Pathogens, however, counteract stomatal immunity by producing effectors and toxins that manipulate stomatal behavior.^4^ For instance, the foliar bacterial pathogen *Pseudomonas syringae* pv. *tomato* DC3000 (*Pto*) secretes coronatine (COR), a mimic of the bioactive phytohormone jasmonic acid-isoleucine (JA-Ile) to suppress stomatal closure.^11^. While COR production is specific to certain *Pseudomonas* species, many foliar pathogens activate JA signaling for stomatal invasion by other means such as effectors.^13–16^ COR activates JA signaling via derepressing the master transcriptional regulator of JA signaling MYC2. Activated MYC2 directly regulates the expression of the transcription factors *ANAC019*, *ANAC055*, and *ANAC072*, which directly suppress the phytohormone salicylic acid (SA) biosynthesis gene *ICS1* and activate SA degradation genes, leading to decreased SA accumulation.^17^ This suggests that SA suppression is required for COR-induced stomatal opening. COR is also known to suppress ABA-induced stomatal closure. While MAMP-induced stomatal closure requires ABA biosynthesis, SA biosynthesis is not essential for ABA-induced stomatal closure, placing ABA downstream of SA in MAMP-induced stomatal closure.^11,18^ Therefore, SA suppression does not fully explain the function of COR in suppressing MAMP-induced stomatal closure.

Most plants exhibit a diurnal pattern of stomatal behavior, opening at dawn to maximize photosynthetic activity and closing at dusk to conserve water.^19–21^ This dawn opening aligns with the timing of pathogen invasion, suggesting that pathogens have adapted to exploit plant physiological processes.^22^ Insufficient stomatal opening restricts CO_2_ uptake, leading to excess energy accumulation in chloroplasts and ROS generation, which damages cellular components such as chlorophyll.^19,23,24^ Thus, plants face a physiological trade-off between gas exchange and defense mechanisms. Recently, it has been shown that the blue-light photoreceptor CRY1 functions in both blue light-induced stomatal opening and MAMP-mediated stomatal closure, balancing stomatal defense and gas exchange.^25^ Nevertheless, despite its significance, the evolutionary diversification of trade-offs at stomata remains unexplored.

In this study, we demonstrate that *CYP707A1*, encoding an ABA degradation enzyme,^26,27^ is essential for suppressing SA-, ABA-, and flg22-induced stomatal closure mediated by COR in *Arabidopsis thaliana*. COR induces *CYP707A1* expression through MYC2 binding to a G-box-like element in the *CYP707A1* promoter, facilitating *Pto* invasion. Intriguingly, we found that *CYP707A1*, along with *MYC2* and the JA receptor gene *COI1*, is required for rapid stomatal opening at dawn. Mutants deficient in *CYP707A1* exhibited increased resistance to *Pto* but showed reduced gas exchange in the morning and chlorophyll contents, representing a tradeoff between defense and a growth process. Comparative analyses across Brassicaceae species revealed that while *A. thaliana* and *Arabidopsis lyrata* exhibit rapid stomatal opening at dawn and susceptibility to stomatal invasion by *Pto*, *Capsella rubella* and *Eutrema salsugineum* show the opposite traits. Promoter swap experiments identified the *CYP707A1* promoter as the determinant of this evolutionary trade-off. Overall, we identify a critical regulator for the stomatal aperture, with its evolutionary diversification driving a trade-off between defense and physiological efficiency.

## RESULTS

### *CYP707A1* is essential for COR-mediated stomatal opening and *Pto* invasion

A previous study demonstrated that SA suppression is required for COR-mediated stomatal opening.^17^ However, inhibition of ABA-induced stomatal closure by COR, combined with the positioning of ABA downstream of SA in MAMP-induced stomatal closure,^11,18^ suggests an additional underlying mechanism. Based on this, we hypothesized that COR suppresses ABA pathways to inhibit MAMP-induced stomatal closure. In *A. thaliana*, ABA is inactivated by ABA 8’-hydroxylases, encoded by *CYP707A1* to *CYP707A4*, into phaseic acid (PA) ^26,28^ (Figure S1A). We first measured the expression levels of *CYP707As* in *A. thaliana* Col-0 wild-type (WT) plants following infection with *Pto* or *Pto cor^−^* (a COR-deficient mutant). RT-qPCR analysis revealed that *Pto*, but not *Pto cor^−^*, induces *CYP707A1* expression. This induction was dependent on the JA receptor *COI1*, consistent with the previous report,^17^ while the other three *CYP707As* were not affected (Figures 1A and S1B). Further analysis showed that COR-induced *CYP707A1* expression requires the JA signaling master regulator MYC2 and its homologs MYC3 and MYC4 (Figure 1B), but COR did not induce the expression of the other *CYP707A* genes (Figure S1C). To evaluate whether *CYP707A* genes are necessary for COR-mediated suppression of stomatal closure, we examined mutants of *CYP707A* genes. COR failed to inhibit flg22-triggered stomatal closure in *cyp707a1*, *coi1*, and *myc2 myc3 myc4* (*myc234*) mutant plants but retained this ability in *cyp707a2*, *cyp707a3*, and *cyp707a4* mutant plants (Figures 1C and S1D). Similarly, COR-mediated inhibition of ABA-induced stomatal closure was dependent on *CYP707A1*, *COI1*, and *MYC2 MYC3 MYC4* but not on the other *CYP707As* (Figures 1D and S1E). These findings suggest that COR suppresses flg22- and ABA-induced stomatal closure by upregulating *CYP707A1* expression via JA signaling.

**Figure 1.**
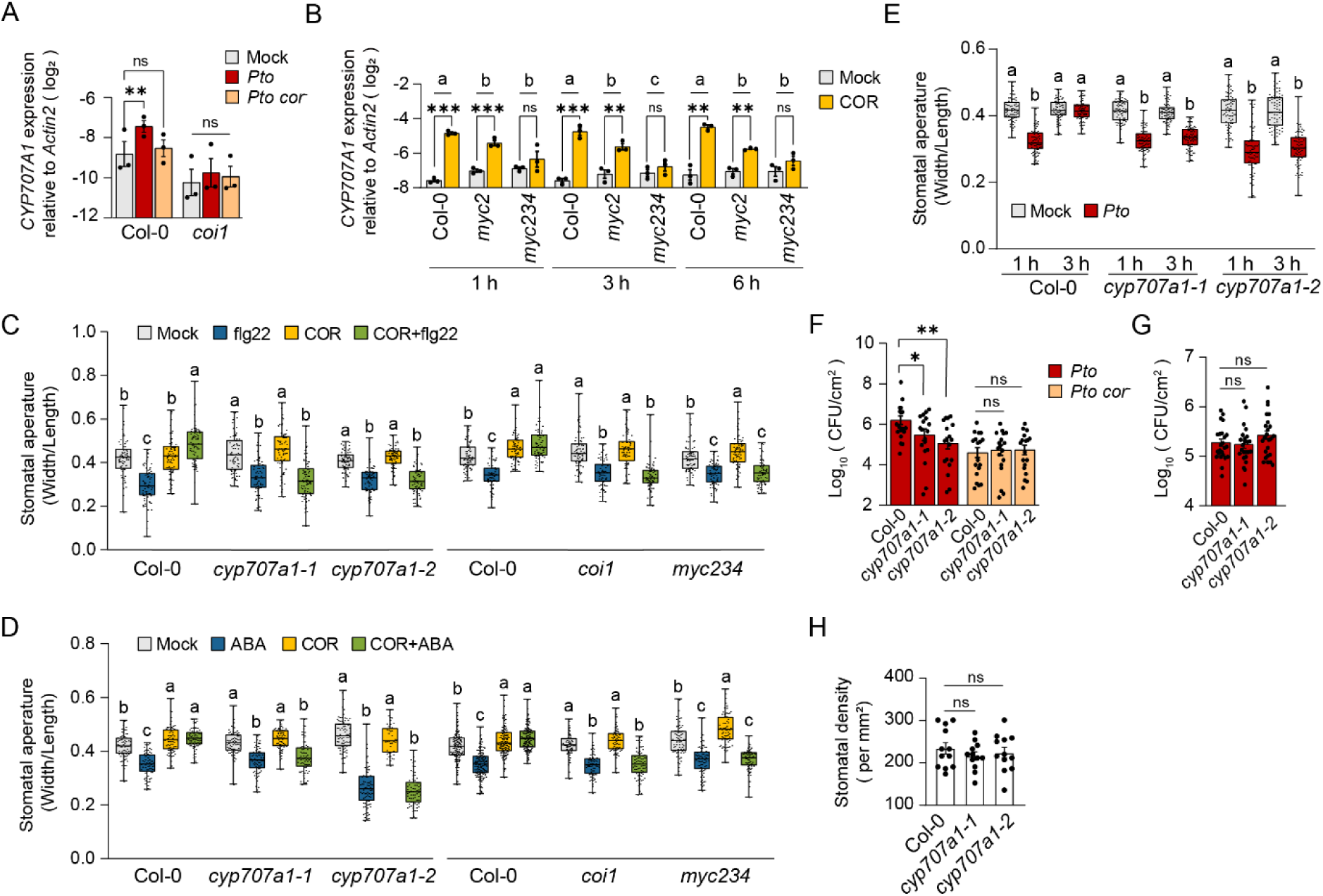
*CYP707A1* is essential for COR-mediated stomatal opening and *Pto* invasion. (A) *CYP707A1* expression. Leaves of 4 to 5-week-old *A. thaliana* Col-0 WT plants were syringe-infiltrated with mock (water), *Pto* (OD600=0.001), or COR-deficient *Pto* mutant *Pto cor*^−^ (OD600=0.001). *CYP707A1* expression was measured at 6 hpi by RT-qPCR. (B) *CYP707A1* expression. Seedlings of 14-day-old *A. thaliana* Col-0 WT and mutant plants were treated with 5 μM COR or mock (0.1% DMSO). *CYP707A1* expression was measured at the indicated time points by RT-qPCR. (A and B) Bars represent means and standard errors of log2 expression levels relative to *ACTIN2* calculated from 3 independent experiments. Asterisks indicate significant differences for the comparisons (\**P <* 0.05, \*\**P <* 0.01, and \*\*\**P* < 0.001, two-tailed Student’s t-tests). Different letters denote significance of differences between groups (one-way ANOVA followed by Tukey test, *P* < 0.05). (C and D) Stomatal aperture. Stomatal apertures of 4 to 5-week-old *A. thaliana* Col-0 WT and mutant plants were measured after treatment with 5 μM COR or mock (0.1% DMSO) for 30 min, followed by incubation with (C) 5 μM flg22 or mock (water) or (D) 10 μM ABA or mock (0.1% ethanol) for 1 hour. (E) Stomatal aperture. Stomatal apertures of 4 to 5-week-old *A. thaliana* Col-0 WT and mutant plants were measured at the indicated time points after the treatment with *Pto* (OD600 = 0.2) or mock (MES buffer). (C to E) n=72 biological replicates per sample from 3 independent experiments. Statistical analysis was performed within the same genotype, and the results were analyzed using one-way ANOVA followed by Tukey test. Different letters denote a statistically significant difference within a genotype (P < 0.05). (F and G) Bacterial growth assay. Leaves of 4 to 5-week-old *A. thaliana* Col-0 WT and mutant plants were (F) sprayed-inoculated (OD600=0.2) or (G) syringe-infiltrated (OD600=0.0002) with *Pto* or *Pto cor*^−^, and bacterial titer was measured at (F) 3 days post inoculation (dpi) or (G) 2 dpi. Bars represent means and standard errors calculated from 2 independent experiments with at least 9 biological replicates per experiment. Asterisks indicate significant differences in the indicated comparisons (\**P <* 0.05, \*\**P <* 0.01, and \*\*\**P* < 0.001, two-tailed Student’s t-tests). (H) Stomatal density. One leaf of 4 to 5-week-old *A. thaliana* Col-0 WT and mutant plants was sampled from each of the 4 plants per genotype. Three microscope photos of each leaf were taken to calculate the average number of stomata as one data point. Bars represent means and standard errors calculated from 3 independent experiments with 4 biological replicates per experiment. Asterisks indicate significant differences in the indicated comparisons (\**P <* 0.05, \*\**P <* 0.01, and \*\*\**P* < 0.001, two-tailed Student’s t-tests). (C, E, D) Results are shown as box plots with boxes displaying the 25th–75th percentiles, the center line indicating the median and whiskers extending to the minimum and maximum values. All individual data points are overlaid.

*Pto* is known to initially induce stomatal closure within one hour, followed by stomatal reopening mediated by COR after several hours.^11^ To determine whether *CYP707A1* is required for this stomatal reopening, we measured stomatal aperture in Col-0 WT and *cyp707a1* mutant plants at one and three hours post-inoculation (hpi). At three hpi, *Pto* was unable to reopen the stomata of *cyp707a1* mutant plants (Figure 1E). Next, to assess the role of *CYP707A1* in susceptibility to *Pto*, we performed both spray- and syringe-inoculation experiments. Spray-inoculation, which relies on stomatal entry, revealed that *cyp707a1* mutants exhibit increased resistance to *Pto* compared to Col-0 WT plants (Figure 1F). In contrast, syringe-inoculation, which bypasses stomatal defenses, showed no significant difference in resistance between *cyp707a1* mutant and Col-0 WT plants (Figure 1G). Importantly, *cyp707a1* mutant plants did not display increased resistance to *Pto cor^−^*. Additionally, stomatal densities in Col-0 WT and *cyp707a1* mutant plants were similar, ruling out differences in stomatal number as a factor in the observed resistance phenotype (Figure 1H). Together, we concluded that *CYP707A1* is a susceptible factor for stomatal defense and is actively exploited by *Pto* through COR.

PA, a product of ABA catabolism by CYP707As, is generally regarded as an inactive derivative of ABA. However, Weng et al. demonstrated that PA can activate ABA signaling by selectively binding to a subset of ABA receptors.^29^ PA is subsequently converted into dihydrophaseic acid (DPA) through the action of PA reductase, encoded by *ABH2* in *A. thaliana*. To investigate whether ABH2 is involved in COR-mediated inhibition of flg22-triggered stomatal closure, we conducted RT-qPCR analyses, which revealed that COR does not induce *ABH2* expression (Figure S2A). Further, functional assays showed that *ABH2* is not required for COR-mediated inhibition of flg22-triggered stomatal closure (Figure S2B). These findings indicate that the conversion of PA to DPA is not necessary for COR action.

### *CYP707A1* is essential for COR-mediated inhibition of SA-induced stomatal closure

A previous study demonstrated that *ANAC019*, *ANAC055*, and *ANAC072* are required for COR-mediated inhibition of stomatal closure by suppressing SA accumulation.^17^ Surprisingly, we found that although these *ANAC* transcription factors are induced by COR,^17^ they are neither required for COR-mediated inhibition of flg22-, *Pto,* or ABA-induced stomatal closure, nor for stomatal reopening (Figures 2A, 2C, and 2D). To resolve this apparent discrepancy, we examined the roles of ABA and SA in flg22-triggered stomatal closure. In our experimental conditions, flg22 failed to induce stomatal closure in *aba2* mutant, which is deficient in ABA biosynthesis,^30^ but successfully closed the stomata of *npr1* and *sid2* mutant plants, deficient in SA perception and biosynthesis,^31,32^ respectively (Figure 2A). This indicates that ABA, but not SA, is required for flg22-triggered stomatal closure under our conditions.

**Figure 2.**
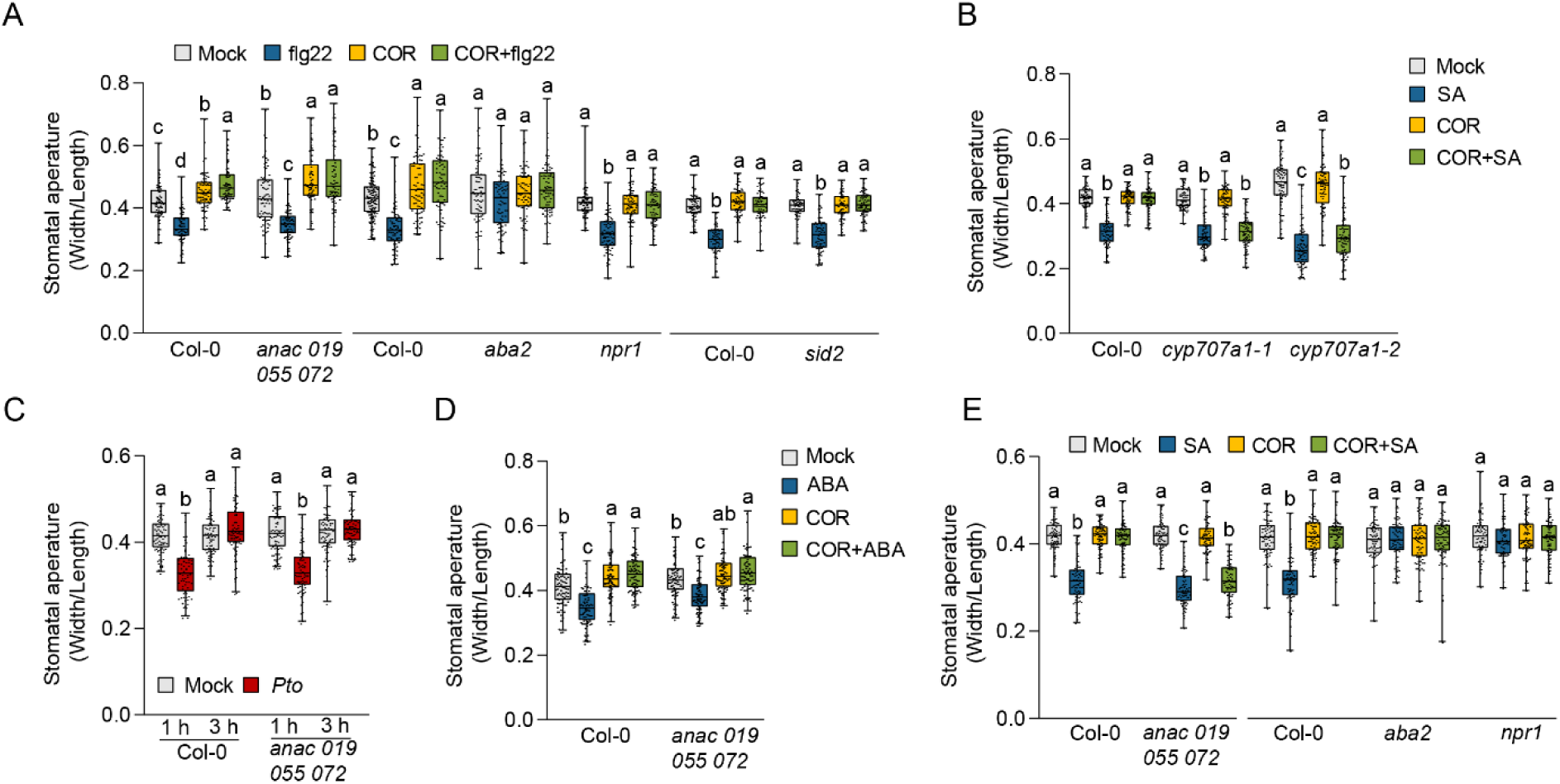
*CYP707A1* is essential for COR-mediated inhibition of SA-induced stomatal closure. (A to E) Stomatal aperture. Stomatal apertures of 4 to 5-week-old *A. thaliana* Col-0 WT and mutant plants were measured after treatment with 5 μM COR or mock (0.1% DMSO) for 30 min, followed by incubation with (A) 5 μM flg22 or mock (water), (D)10 μM ABA or mock (0.1% ethanol), or (B and E) 100 μM SA or mock (water) for 1 hour. (C) Stomatal apertures of 4 to 5-week-old *A. thaliana* Col-0 WT and mutant plants were measured at the indicated time points after the treatment with *Pto* (OD600 = 0.2) or mock (MES buffer). n=72 biological replicates per sample from 3 independent experiments. Results are shown as box plots with boxes displaying the 25th–75th percentiles, the center line indicating the median and whiskers extending to the minimum and maximum values. All individual data points are overlaid. Statistical analysis was performed within the same genotype and analyzed using one-way ANOVA followed by Tukey test. Different letters denote a statistically significant difference within a genotype (*P* < 0.05).

SA, however, is also known to induce stomatal closure.^18^ As expected, SA-induced stomatal closure depended on both *ABA2* and *NPR1* (Figure 2E), demonstrating that ABA accumulation and SA signaling are required for SA-induced stomatal closure. Notably, we observed that *ANAC019*, *ANAC055*, and *ANAC072* are required for COR-mediated inhibition of SA-induced stomatal closure (Figure 2E). These findings suggest that these *ANACs* are required for COR-mediated inhibition of stomatal closure only when SA plays a significant role. Interestingly, *CYP707A1* was also found to be essential for COR-mediated inhibition of SA-induced stomatal closure (Figure 2B). Furthermore, COR still induced *CYP707A1* expression in *anac019 anac055 anac072* (*anac*) mutant plants, indicating that *CYP707A1* functions independently of these ANAC transcription factors (Figure S3). Collectively, these results highlight the indispensable role of *CYP707A1* in COR-mediated inhibition of stomatal closure, regardless of whether it is triggered by flg22, ABA, or SA, underscoring its critical function in COR-mediated stomatal reopening.

### COR fails to open stomata in *C. rubella* and *E. salsugineum*

To explore the evolutionary conservation of COR-mediated stomatal opening within Brassicaceae plants, we selected *A. lyrata*, *C. rubella*, and *E. salsugineum*, which are related to *A. thaliana* to varying extents and have available genome sequences with high-quality gene annotations^33^ (Figure 3A). Phylogenetic analysis revealed that each of these Brassicaceae species possesses four *CYP707A* genes, corresponding to the *A. thaliana CYP707A1, CYP707A2, CYP707A3,* and *CYP707A4* genes (Figure S1A). We then investigated whether COR induces *CYP707A1* expression in these species. In *A. lyrata*, COR induced only *CYP707A1*, similar to *A. thaliana* (Figures 3B and S4B). However, COR failed to induce any of the *CYP707As* genes in *C. rubella* and *E. salsugineum* (Figures 3C, S4C and S4D), although it did induce the JA marker gene *HAI1*^34^ (Figure S4A). This suggests that while *C. rubella* and *E. salsugineum* are capable of sensing COR, they do not induce the *CYP707A1* gene. These findings indicate an evolutionary diversification of the COI1-MYC2-CYP707A1 module within Brassicaceae species. Consistent with these findings, COR inhibited flg22-triggered stomatal closure in *A. thaliana* and *A. lyrata*, but not in *C. rubella* and *E. salsugineum* (Figure 3D).

**Figure 3.**
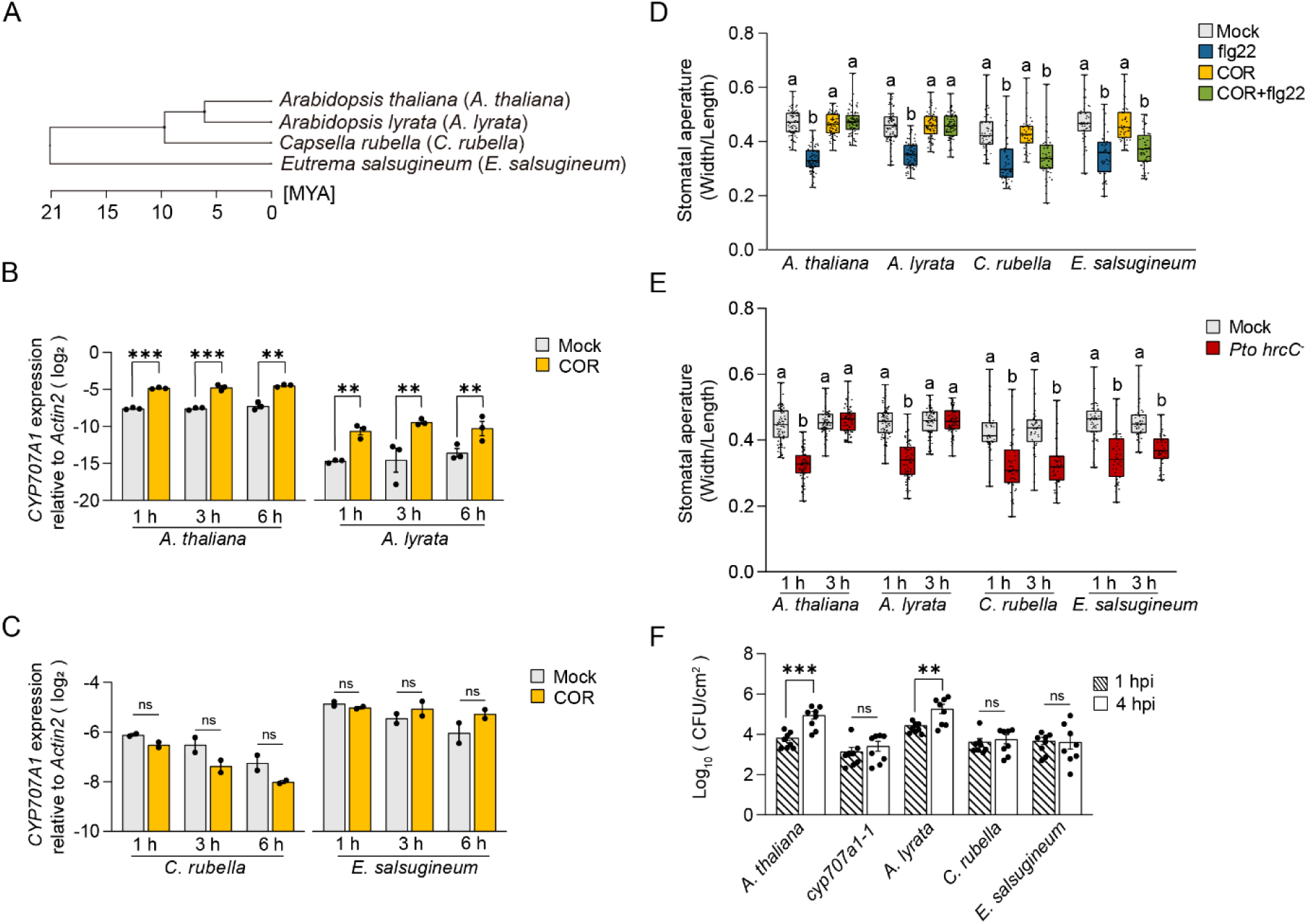
COR fails to open stomata in *C. rubella* and *E. salsugineum*. (A) A phylogenetic tree of the 4 Brassicaceae species used in this study. *A. thaliana* (Col-0); *A. lyrata* (MN47); *C. rubella* (N22697); *E. salsugineum* (Shandong). (B and C) *CYP707A1* expression. Seedlings of 14-day-old Brassicaceae plants were treated with 5 μM COR or mock (0.1% DMSO). *CYP707A1* expression was measured at the indicated time points by RT-qPCR. Bars represent means and standard errors of log2 expression levels relative to *ACTIN2* calculated from 3 (B) or 2 (C) independent experiments. Asterisks indicate significant differences from mock (\**P <* 0.05, \*\**P <* 0.01, and \*\*\**P* < 0.001, Student’s t-tests). (D and E) Stomatal aperture. (D) Stomatal apertures of 4 to 5-week-old Brassicaceae plants were measured after treatment with 5 μM COR or mock (0.1% DMSO) for 30 min, followed by incubation with 5 μM flg22 or mock (water) for 1 hour. (E) Stomatal apertures of 4 to 5- week-old Brassicaceae plants were measured at the indicated time points after the treatment with *Pto hrcC^−^* (OD600 = 0.2) or mock (MES buffer). (D and E) n=72 (*A. thaliana* and *A. lyrata*); n=44 (*C. rubella* and *E. salsugineum*) biological replicates per sample from at least 2 independent experiments. Results are shown as box plots with boxes displaying the 25th–75th percentiles, the center line indicating the median and whiskers extending to the minimum and maximum values. All individual data points are overlaid. Statistical analysis was performed within the same genotype, and the results were analyzed using one-way ANOVA followed by Tukey test. Different letters denote a statistically significant difference within a genotype (P < 0.05). (F) *Pto* invasion assay. Detached leaves of 4 to 5-week-old Brassicaceae plants were soak-inoculated with *Pto hrcC^−^* suspension (OD600 = 0.2). Endophytic bacterial titer was determined at 1 and 4 hpi. Experiments were repeated at least 2 times each with 8 biological replicates with similar results. Asterisks indicate significant differences from 1 hpi (\**P <* 0.05, \*\**P <* 0.01, and \*\*\**P* < 0.001, two-tailed Student’s t-tests).

Next, we investigated whether *Pto* could induce stomatal reopening in these Brassicaceae species. We previously showed that *Pto* appears to activate effector-triggered immunity in *E. salsugineum*, which may counteract COR-mediated stomatal reopening.^35^ To address this, we used *Pto hrcC^−^*, a mutant that lacks the functional type III secretion system but retains COR production.^36^ Our results showed that at one hpi, *Pto hrcC^−^* triggered stomatal closure in all tested species, similar to *Pto* (Figure 3E). However, during stomatal reopening, we observed species-specific differences. While *Pto hrcC^−^*was able to reopen stomata in *A. thaliana* and *A. lyrata*, it failed to do so in *C. rubella* and *E. salsugineum* (Figure 3E). To more directly assess the role of *CYP707A1* in stomatal defense, we performed *Pto* stomatal entry assays.^37^ Detached leaves were treated with *Pto hrcC^−^*, and its endophytic population was measured at one and four hpi, during which no bacterial multiplication occurred. The results revealed significant *Pto hrcC^−^* invasion into the leaf interior of *A. thaliana* and *A. lyrata* between one and four hpi (Figure 3F). However, consistent with the inability of COR to reopen stomata in *C. rubella* and *E. salsugineum*, *Pto hrcC^−^* invasion was hardly observed in these species (Figure 3F). Thus, COR effectively mediates stomatal reopening in *A. thaliana* and *A. lyrata*, but fails to do so in *C. rubella* and *E. salsugineum*, further highlighting the evolutionary diversification of the COI1-MYC2-CYP707A1 module within Brassicaceae species.

### The promoter of *CYP707A1* underlies the evolutionary diversification of the COI1-MYC2-CYP707A1 module

To investigate the factors underlying the evolutionary diversification of COR action, we hypothesized that differences in *CYP707A1* genes might be responsible, given that COR can induce the JA marker gene in *C. rubella* and *E. salsugineum* (Figure S4A). To test this hypothesis, we performed promoter-swap experiments. We generated transgenic *A. thaliana cyp707a1* mutants and *E. salsugineum* plants expressing *A. thaliana* or *E. salsugineum CYP707A1* coding sequences (CDS) under the *CYP707A1* promoter from *A. thaliana*, *C. rubella*, or *E. salsugineum*, in various combinations (Figure S5A). Our results showed that flg22 consistently triggered stomatal closure in all transgenic lines (Figures 4A, 4C, S5B, and S5C). However, COR inhibited flg22-triggered stomatal closure in *A. thaliana cyp707a1* mutant and *E. salsugineum* plants carrying any of the tested *CYP707A1* CDSs regulated by the *A. thaliana CYP707A1* promoter (Figures 4A, 4C, S5B, and S5C). This finding suggests that the *CYP707A1* promoter sequence is the key determinant in the evolutionary diversification of the COI1-MYC2-CYP707A1 module, which governs COR-mediated stomatal opening within the Brassicaceae family.

**Figure 4.**
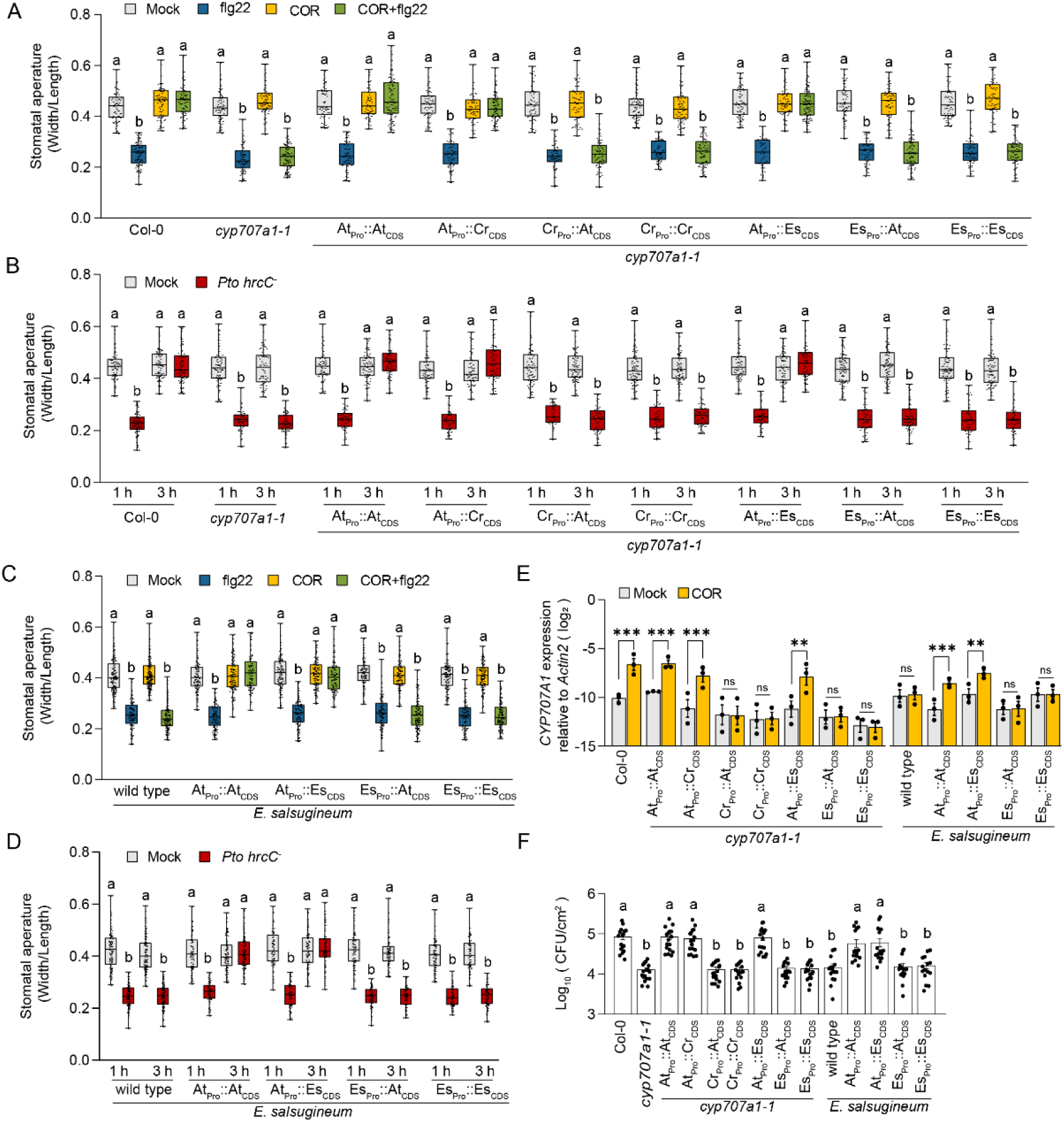
The *CYP707A1* promoter underlies the evolutionary diversification of the COI1-MYC2-CYP707A1 pathway. (A to D) Stomatal aperture. Stomatal apertures of 4 to 5-week-old *A. thaliana* (A and B) and *E. salsugineum* (C and D) plants were measured after treatment with (A and C) 5 μM COR or mock (0.1% DMSO) for 30 min, followed by incubation with 5 μM flg22 and mock (water); (B and D) *Pto hrcC^−^* (OD600 = 0.2) or mock (MES buffer) at the indicated time points. n≥50 biological replicates per sample from 3 independent experiments. Results are shown as box plots with boxes displaying the 25th–75th percentiles, the center line indicating the median and whiskers extending to the minimum and maximum values. All individual data points are overlaid. Statistical analysis was performed within the same genotype, and the results were analyzed using one-way ANOVA followed by Tukey test. Different letters denote a statistically significant difference within a genotype (P < 0.05). (E) *CYP707A1* expression. Seedlings of 14-day-old *A. thaliana* Col-0 WT and mutant plants were treated with 5 μM COR or mock (0.1% DMSO). *CYP707A1* expression was measured at 6 hpi by RT-qPCR. Bars represent means and standard errors of log2 expression levels relative to *ACTIN2* calculated from 3 independent experiments. Asterisks indicate significant differences from mock (\**P <* 0.05, \*\**P <* 0.01, and \*\*\**P* < 0.001, two-tailed Student’s t-tests). (F) *Pto* invasion assay. Detached leaves of 4 to 5-week-old *A. thaliana* and *E. salsugineum* plants were soak-inoculated with *Pto hrcC^−^* suspension (OD600 = 0.2). Endophytic bacterial titer was determined at 3 hpi. Bars represent means and standard errors from 3 independent experiments each with at least 5 biological replicates. The results were analyzed using one-way ANOVA followed by Tukey test. Different letters denote a statistically significant difference (P < 0.05).

We then examined whether *Pto hrcC^−^* could induce stomatal reopening in these transgenic plants. Consistent with COR-mediated inhibition of flg22-triggered stomatal closure, *Pto hrcC^−^* reopened stomata in *A. thaliana cyp707a1* and *E. salsugineum* plants carrying any of *CYP707A1* CDSs under the regulation of the *A. thaliana CYP707A1* promoter. In contrast, *Pto hrcC^−^* failed to reopen stomata in transgenic plants with any of *CYP707A1* CDS driven by the *C. rubella* or *E. salsugineum* promoters (Figures 4B, 4D, S5D, and S5E). To further corroborate these findings, we examined *CYP707A1* expression in these transgenic plants. We found that COR induces *CYP707A1* expression only in plants with the *A. thaliana CYP707A1* promoter, not in those with the *C. rubella* or *E. salsugineum* promoter, which correlated with the observed stomatal responses (Figure 4E).

Finally, we conducted *Pto* stomatal invasion assays in these transgenic plants. Plants carrying *CYP707A1* driven by the *C. rubella* or *E. salsugineum* promoter, as well as *cyp707a1* mutant plants, exhibited greater resistance to *Pto hrcC^−^* invasion, compared to Col-0 WT and transgenic plants with *CYP707A1* driven by the *A. thaliana* promoter (Figure 4F). Together, these results further demonstrated that the *CYP707A1* promoter is the key determinant in the evolutionary diversification, regulating COR-mediated *Pto* virulence within the Brassicaceae family.

### MYC2 directly binds to the *CYP707A1* promoter to drive COR-induced *CYP707A1* expression

The dependency of COR-induced *CYP707A1* expression on *MYC2 MYC3 MYC4* suggests that MYC2 directly regulates *CYP707A1* transcription (Figure 1B). To investigate this, we analyzed the sequence of the *A. thaliana CYP707A1* promoter and identified one canonical G-box motif—a known MYC2 binding site—and two G-box-like motifs within 1000 bp upstream of the *CYP707A1* start codon (Figure 5A). To investigate MYC2 binding, we performed chromatin immunoprecipitation (ChIP) assays using transgenic *A. thaliana* plants expressing MYC2-Flag fusion protein driven under the control of its native promoter. We designed primers targeting the *CYP707A1* promoter region, containing the G-box, G-box-like 1, or G-box-like 2 motif (Figure 5A). The promoter region of *ANAC019*, a previously established MYC2 target,^17^ was used as a positive control, and the *eIF4A* and the *CYP707A1* CDS served as the reference and a negative control, respectively. ChIP analysis revealed significant enrichment of MYC2 at the G-box, the G-box-like 1, and the G-box-like 2, regardless of COR treatment (Figure 5B). These results suggest that MYC2 directly regulates *CYP707A1* expression through its interaction with these motifs.

**Figure 5.**
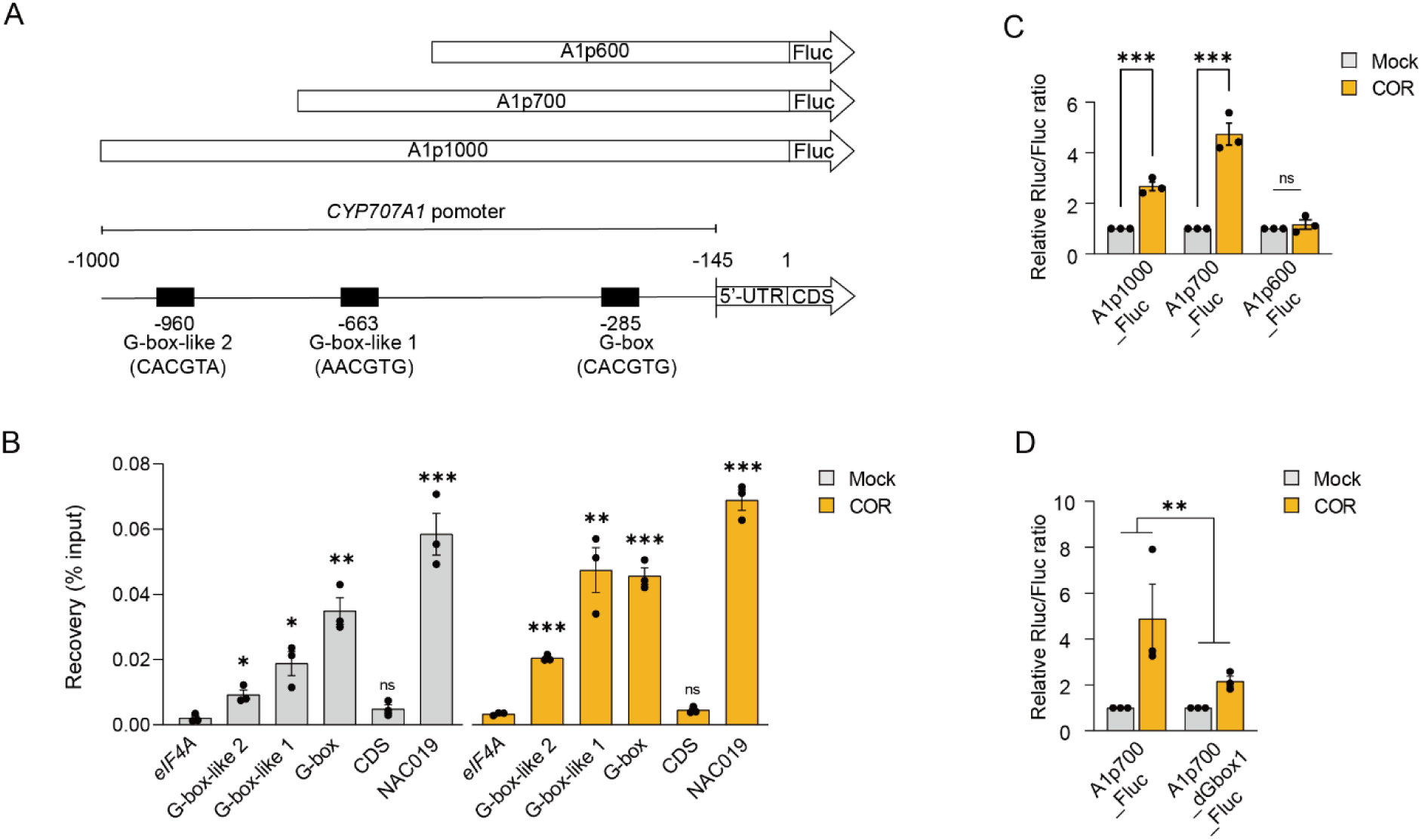
MYC2 directly regulates *CYP707A1* expression through its interaction with the G-box-like 1 motif. (A) Schematic diagram of the *A. thaliana CYP707A1* promoter. The G-box (CACGTG), along with the G-box-like 1 (AACGTG) and G-box-like 2 (CACGTA) motifs, are positioned 285 bp, 663 bp, and 960 bp upstream of the start codon, respectively. CDS is the coding sequence of *CYP707A1*. 5’ UTR is the 5’ untranslated region. (B) ChIP assay. Seedlings of 14-day-old Col-0 and pMYC2::MYC2-Flag plants were treated with 5 μM COR or mock (0.1% DMSO) and harvested at 3 h. Bars represent means and standard errors of the percentage of input values of the ChIP DNA, calculated from 3 independent experiments. Asterisks indicate statistically significant differences from *eIF4A* within the respective treatment (*P<0.05, **P<0.01; ***P<0.001, two-tailed Student’s t-tests). (C and D) *Agrobacterium* strains harboring Firefly luciferase (Fluc) reporter constructs together with *Agrobacterium* strain carrying the control Renilla luciferase construct (Rluc) were mixed at a 10:1 ratio (final OD600 of 0.1) and infiltrated into leaves of 3 to 4-week-old *A. thaliana dde2 ein2 pad4 sid2* mutant plants. After two days, the infiltrated leaves were treated with 5 µM COR for 3 hours. Relative Fluc/Rluc values are shown (mock set to 1). Bars represent means and standard errors calculated from 3 independent experiments. Asterisks indicate significant differences in the indicated comparisons (\**P <* 0.05, \*\**P <* 0.01, and \*\*\**P* < 0.001, two-tailed Student’s t-tests).

To evaluate the functional contribution of these motifs to *CYP707A1* promoter activity, we constructed a series of *CYP707A1* promoter-luciferase reporter vectors: A1p1000_Fluc (containing G-box, G-box-like 1, and G-box-like 2), A1p700_Fluc (containing G-box and G-box-like 1), and A1p600_Fluc (containing only G-box). These constructs were used in dual-luciferase reporter assays via *Agrobacterium*-mediated transient expression in *A. thaliana dde2 ein2 pad4 sid2* mutant plants, which enables efficient *Agrobacterium*-mediated transient transformation in *A. thaliana.*^38^ The reporter assay revealed significant COR-induced luciferase activity in plants harboring A1p1000_Fluc and A1p700_Fluc, with A1p700_Fluc showing the strongest induction (Figure 5C). In contrast, no induction was observed in plants with A1p600_Fluc, indicating that the G-box-like 1 motif is essential for COR responsiveness (Figure 5C). To further confirm the role of the G-box-like 1 motif, we generated a mutant version, A1p700_dGbox1_Fluc, with a disrupted G-box-like 1 motif. Dual-luciferase assays demonstrated that this mutation significantly attenuated COR-induced reporter activity (Figure 5D). These findings suggest that MYC2 directly binds to at least the G-box-like 1 motif in the *CYP707A1* promoter, driving COR-induced *CYP707A1* expression in *A. thaliana*.

In our analysis of over 1,000 *A. thaliana* accessions,^39^ we identified the G-box-like 1 motif within the *CYP707A1* promoter region at the corresponding positions in the remaining 780 accessions, after excluding accessions with unreadable sequences. This strong conservation highlights the essential role of this motif in regulating *CYP707A1* expression across *A. thaliana* populations. Interestingly, G-box-like 1 is also conserved in *A. lyrata*, *C. rubella*, and *E. salsugineum*, suggesting that while necessary for COR-induced *CYP707A1* expression, it alone is not sufficient. However, the highly divergent promoter sequences of *CYP707A1* among the Brassicaceae species hindered the identification of specific elements responsible for the evolutionary diversification of *CYP707A1* regulation.

### *CYP707A1* is required for rapid stomatal opening at dawn

The evolutionary diversification of *CYP707A1* regulation, which conferred resistance to COR-mediated *Pto* invasion in *C. rubella* and *E. salsugineum* raises the question of why *A. thaliana* and *A. lyrata* retained this regulatory mechanism despite its negative consequence in pathogen resistance. Given the regulation of JA signaling by the circadian clock,^40^ and the established role of light in modulating stomatal aperture, we hypothesized that the COI1-MYC2-CYP707A1 module might play a role in light-mediated stomatal responses. To test this hypothesis, we examined *CYP707A1* expression in response to light. Our analysis revealed a rapid induction of *CYP707A1* expression within one hour of light exposure (Figures 6A and 6B). To further elucidate the role of the COI1-MYC2-CYP707A1 module in stomatal responses to light, we conducted time-course experiments assessing stomatal aperture before and after light exposure in *A. thaliana* Col-0 WT plants and *cyp707a1*, *coi1*, and *myc234* mutant plants (Figure 6C). In Col-0 WT plants, we observed rapid stomatal opening within one hour of light exposure. However, this response was significantly delayed in *cyp707a1*, *coi1*, and *myc234* mutant plants (Figure 6C). These findings underscore the involvement of JA signaling in light-mediated stomatal responses and highlight the critical role of the COI1-MYC2-CYP707A1 module in coordinating rapid stomatal opening at dawn.

**Figure 6.**
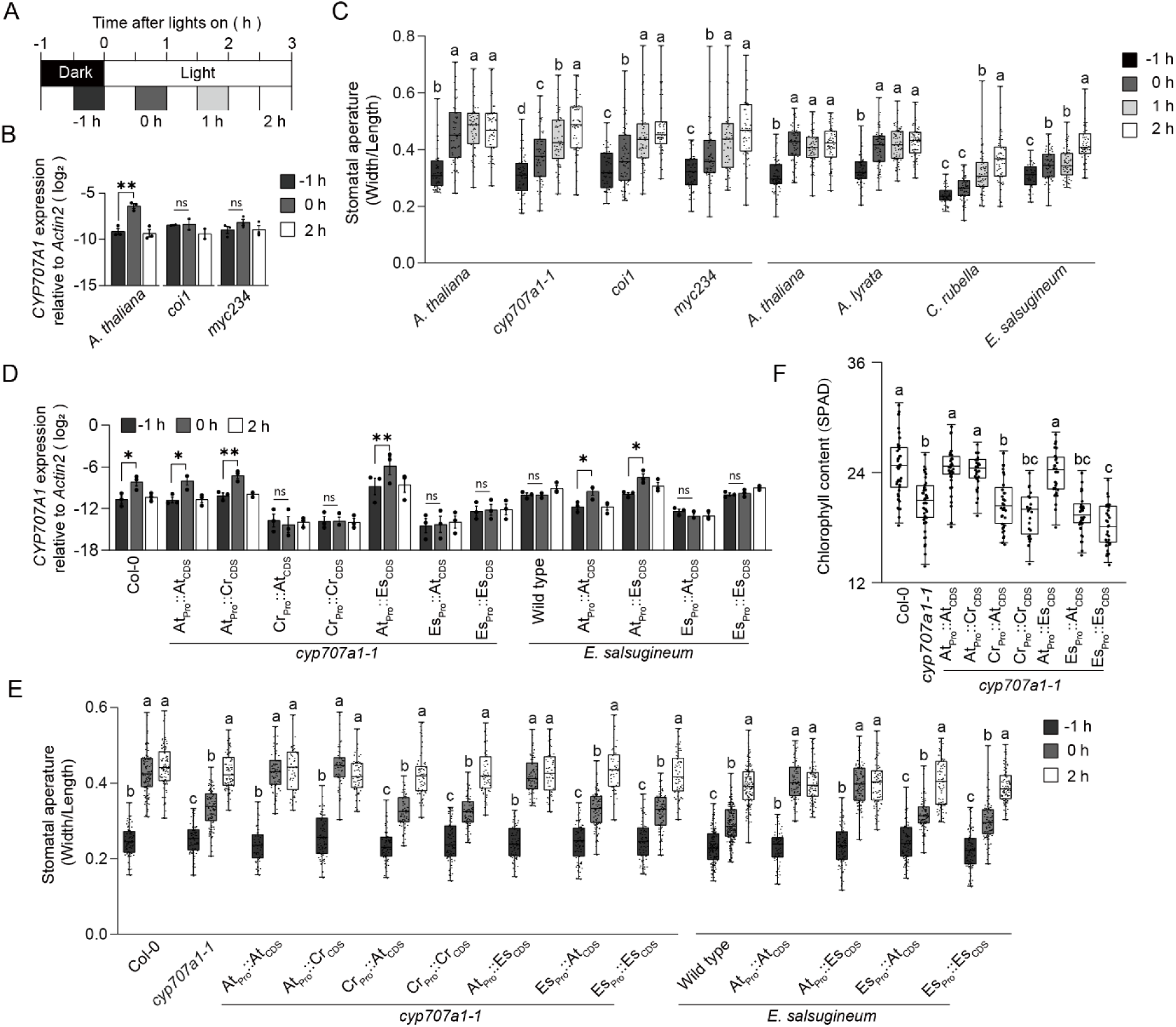
*CYP707A1* regulates light-induced stomata opening at dawn. (A) The time frame of sampling. (B) *CYP707A1* expression. Leaves of 4 to 5-week-old *A. thaliana* Col-0 WT and mutant plants were harvested at the indicated time points, and *CYP707A1* expression was measured by RT-qPCR. Bars represent means and standard errors of log2 expression levels relative to *ACTIN2* calculated from three independent or two (*coi1*) experiments. Asterisks indicate significant differences for the indicated comparison (\**P <* 0.05, \*\**P <* 0.01, and \*\*\**P* < 0.001, two-tailed Student’s t-test). (C) Stomatal aperture. Stomatal apertures of 4 to 5-week-old *A. thaliana* Col-0 WT and mutant and other Brassicaceae plants were measured at the indicated time points. n≥60 for Col-0, *cyp707a1-1*, *coi1*, and *myc234*; n≥60 for *A. lyrata*; n≥40 for *C. rubella* and *E. salsugineum*. Statistical analysis was performed within the same genotype, and the results were analyzed using one-way ANOVA followed by Tukey test. Different letters denote a statistically significant difference within a genotype (P < 0.05). (D) *CYP707A1* expression. Leaves of 4 to 5-week-old plants were harvested at the indicated time points, and *CYP707A1* expression was measured by RT-qPCR. Bars represent means and standard errors of log2 expression levels relative to *ACTIN2* calculated from three independent experiments. Asterisks indicate significant differences for the indicated comparisons (\**P <* 0.05, \*\**P <* 0.01, and \*\*\**P* < 0.001, two-way ANOVA followed by Tukey test). (E) Stomatal aperture. Stomatal apertures of 4 to 5-week-old plants were measured at the indicated time points. n≥50 for each sample from 3 independent experiments. Statistical analysis was performed within the same genotype, and the results were analyzed using one-way ANOVA followed by Tukey test. Different letters denote a statistically significant difference within a genotype (P < 0.05). (F) Chlorophyll content. Chlorophyll content in leaves of 4 to 5-week-old plants was measured by SPAD. n≥18 for each sample from 3 independent each with at least 6 biological replicates. The results were analyzed using one-way ANOVA followed by Tukey test. (C, E, F) Results are shown as box plots with boxes displaying the 25th–75th percentiles, the center line indicating the median, and whiskers extending to the minimum and maximum values. All individual data points are overlaid.

### *CYP707A1* promoter underlies delayed stomatal opening at dawn in *C. rubella* and *E. salsugineum*

To investigate whether the sensitivity to COR-mediated stomatal reopening is evolutionarily linked to light-mediated stomatal opening, we examined Brassicaceae species under light exposure. Consistent with their sensitivity to COR, *A. thaliana* and *A. lyrata*, which exhibit COR-mediated stomatal reopening by *Pto*, showed rapid stomatal opening at dawn (Figure 6C). In contrast, *C. rubella* and *E. salsugineum*, which do not exhibit COR-mediated stomatal reopening, displayed delayed stomatal opening, resembling the phenotypes of *coi1*, *myc234*, and *cyp707a1* mutant *A. thaliana* plants (Figure 6C). This suggests a link between the absence of COR-mediated stomatal reopening and delayed stomatal opening at dawn.

To determine whether the *CYP707A1* promoter governs rapid stomatal opening at dawn, we analyzed *CYP707A1* expression and stomatal aperture in promoter-swap transgenic lines. In these lines, *CYP707A1* driven by the *A. thaliana CYP707A1* promoter was rapidly induced within one hour of light exposure, whereas *CYP707A1* driven by the *C. rubella* or *E. salsugineum CYP707A1* promoter showed no such induction (Figure 6D). Stomatal aperture analysis revealed that *A. thaliana cyp707a1* and *E. salsugineum* plants containing *A. thaliana* promoter-driven *CYP707A1* exhibited rapid stomatal opening at dawn akin to WT *A. thaliana*, while plants with other promoter constructs did not (Figures 6E and S6A). These results indicate that the *A. thaliana CYP707A1* promoter is pivotal for rapid stomatal opening at dawn and further links the COR-mediated suppression of flg22-triggered stomatal closure, COR-mediated stomatal reopening, and light-induced rapid stomatal opening via a shared COI1-MYC2-CYP707A1 signaling module.

To further assess the functional significance of this module, we measured stomatal conductance and transpiration rates during the dark-to-light transition at dawn. Plants carrying *CYP707A1* driven by the *A. thaliana* promoter exhibited higher stomatal conductance and transpiration rates in the morning compared to plants with *CYP707A1* driven by the *C. rubella* or *E. salsugineum* promoter (Figures S6B and S6C). This reduced stomatal conductance likely affects physiological traits, such as chlorophyll accumulation, particularly under high-light conditions. To test this, plants grown under normal light (200 μmol m^−2^ s^−1^) were exposed to high light (600 μmol m^−2^ s^−1^), and chlorophyll content was measured after two days. Plants expressing any of *CYP707A1* CDSs driven by the *A. thaliana* promoter maintained normal chlorophyll levels, whereas those with *CYP707A1* CDSs driven by the *C. rubella* or *E. salsugineum* promoter showed significant chlorophyll reductions, similar to *A. thaliana cyp707a1* mutant plants (Figure 6F). Altogether, our findings suggest that the *CYP707A1* promoter acts as an evolutionary pivot, regulating the stomatal response to light and pathogen virulence across Brassicaceae species. This regulation represents a trade-off between pathogen resistance and physiological efficiency, providing insights into how evolutionary pressures shape the balance between plant immunity and growth.

## DISCUSSION

This study uncovers a critical role for *CYP707A1* in mediating the evolutionary trade-off between stomatal defense and gas exchange in Brassicaceae plants. By demonstrating that *CYP707A1* expression is essential for COR-mediated inhibition of stomatal closure and rapid stomatal opening at dawn, we underscore its dual role in pathogen susceptibility and physiological efficiency. The regulatory diversification of the *CYP707A1* promoter among Brassicaceae species reflects an evolutionary balance between these competing demands, offering insights into how transcriptional regulation optimizes growth-defense trade-offs in plants (Figure 7).

**Figure 7.**
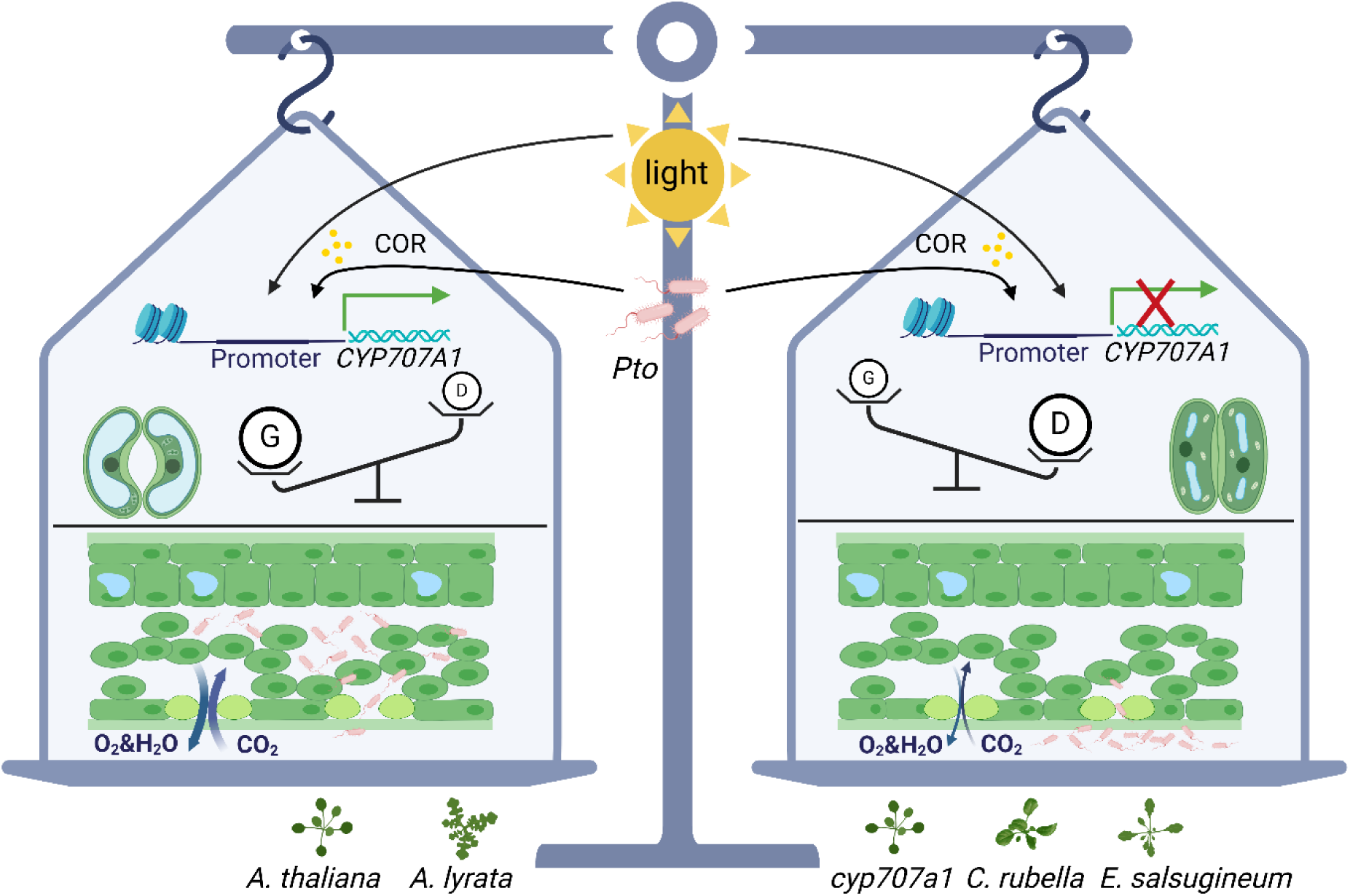
Model illustrating the evolutionary trade-off between stomatal defense and gas exchange. In *A. thaliana* and *A. lyrata*, *Pto* exploits the JA-mediated induction of *CYP707A1* by COR to promote stomatal opening, facilitating bacterial invasion. Notably, this JA-*CYP707A1* pathway also regulates rapid stomatal opening at dawn, a process critical for efficient gas exchange. This dual role highlights an evolutionary trade-off between defense mechanisms and growth-related physiological processes. Conversely, in *cyp707a1* mutant, *C. rubella* and *E. salsugineum*, the JA pathway is decoupled from *CYP707A1* regulation, conferring resistance to *Pto* invasion and leading to delayed stomatal opening at dawn. These evolutionary trade-offs are driven by diversification in the *CYP707A1* promoter regions. Abbreviations: *Pto* (*Pseudomonas syringae pv. tomato* DC3000), COR (Coronatine). D (Defense), G (Gas exchange). This illustration was created with BioRender.com.

*CYP707A1* operates at the intersection of JA and ABA signaling pathways, as evidenced by its induction through MYC2 binding to specific promoter motifs. This mechanism allows pathogens such as *P. syringae* to exploit stomatal responses for invasion. Concurrently, *CYP707A1*-driven stomatal opening enhances gas exchange and maintains chlorophyll production under high light, exemplifying a physiological trade-off. Promoter-swap experiments confirm that differences in *CYP707A1* promoter activity, rather than variations in the CYP707A1 protein or MYC2 transcription factor, dictate the evolutionary trade-offs observed across Brassicaceae species. This divergence explains why *A. thaliana* and *A. lyrata* exhibit rapid stomatal opening at dawn and increased COR-induced susceptibility, whereas *C. rubella* and *E. salsugineum* show delayed stomatal opening and heightened resistance to pathogen invasion. These findings suggest that promoter evolution drives the observed functional divergence, with species prioritizing either pathogen resistance or gas exchange efficiency based on ecological pressures. Consistent with this, previous work indicates that a shift in transcriptional start site appears to mediate an evolutionary trade-off in SA accumulation within Brassicaceae,^41^ underscoring the broader significance of promoter evolution in shaping plant traits.

Research has demonstrated that *Pto* reopens stomata in tomato plants through COR activity.^42^ Although the exact mechanism of COR-mediated stomatal opening in tomatoes remains unclear, the presence of two *CYP707A* genes in the tomato genome homologous to the Brassicaceae *CYP707A1/A3* branch might suggest a conserved mechanism. If true, *CYP707A1*-mediated stomatal opening may represent an ancestral function that has been lost in *C. rubella* and *E. salsugineum* through evolutionary modifications in their promoters. Alternatively, this trait may have evolved in the common ancestor of *A. thaliana* and *A. lyrata* through promoter diversification. Regardless of whether this reflects an evolutionary gain or loss, the diversification of *CYP707A1* regulation highlights a trade-off balancing pathogen resistance and gas exchange efficiency. These findings refine existing models of the growth-defense trade-off by emphasizing the integration of light-responsive stomatal dynamics with pathogen susceptibility.

Stomatal closure is governed by a wide range of internal and external stimuli. In addition to ABA and SA, other phytohormones such as brassinosteroid and ethylene have been shown to induce stomatal closure.^43,44^ Notably, exogenous JA can also trigger stomatal closure under certain conditions.^45,46^ Our findings indicate that flg22- and SA-triggered stomatal closure depends on ABA accumulation, although prior research indicates that MAMPs can also induce stomatal closure via ABA-independent pathways to some extent.^47^ Additionally, ABA is not essential for dak-induced stomatal closure but determines response speed.^48^ These results illustrate the diversity of mechanisms plants employ to regulate stomatal closure. Importantly, *CYP707A1* plays a pivotal role in COR-induced inhibition of flg22-, ABA-, and SA-triggered stomatal closure while promoting *Pto*-mediated stomatal reopening and facilitating rapid stomatal opening at dawn. This underscores the critical role of suppressing ABA accumulation in enabling effective stomatal opening, even as plants rely on multiple pathways for closure. A recent study reveals that COR promotes the nuclear transport of a negative regulator of ABA signaling, effectively antagonizing ABA’s influence and reopening stomata.^49^ This dual suppression of ABA accumulation and signaling by COR appears to be fundamental to its function. The identification of the COI1-JA-CYP707A1 pathway as integral to COR-mediated stomatal dynamics and dawn-associated rapid stomatal opening provides a robust foundation for exploring regulatory networks that balance plant defense and physiological efficiency.

While this study identifies the *CYP707A1* promoter as the key determinant of stomatal opening mediated by the COI1-MYC2-CYP707A1 pathway, the exact evolutionary mechanisms driving promoter diversification remain unresolved. Future research should investigate the role of epigenetic modifications, such as DNA methylation, in shaping promoter activity, and conduct fine mapping of DNA sequences governing distinct COR-mediated *CYP707A1* regulation. Broader comparative analyses across Brassicaceae and other plant lineages could reveal the prevalence of these trade-offs and whether similar mechanisms operate in different ecological contexts. Moreover, future research into the interplay between *CYP707A1* regulation and abiotic stress responses, such as drought, would provide a more comprehensive understanding of its role in plant adaptation. Such insights could ultimately inform strategies to leverage these mechanisms for enhancing crop resilience and sustainability in agriculture.

## STAR★METHODS

- KEY RESOURCES TABLE
- RESOURCE AVAILABILITY

- Lead contact
- Materials availability
- EXPERIMENTAL MODEL AND STUDY PARTICIPANT DETAILS
- METHOD DETAILS

- Plant materials and growth conditions
- Bacterial cultivation
- Bacterial growth assay
- Bacterial invasion assay
- RT-qPCR
- ChIP-qPCR analysis
- Luciferase reporter assay
- Stomatal aperture measurement and analysis
- Stomatal conductance and transpiration rate analysis
- Stomatal density analysis
- Chlorophyll content detection
- Molecular cloning
- Phylogenetic tree construction
- Media and reagents
- Statistical analyses

## ACKNOWLEDGEMENTS

This work was supported by the National Natural Science Foundation of China (32170298 and 32250710139 to K.T.), the National Key R&D Program of China (2022YFA1304400 to K.T. and X.H.), JSPS KAKENHI (21H05151, 24K01757, and 22K19178 to A.M., 24KJ0131 to R.H.), and JST PRESTO (JPMJPR17Q6 to A.M.) and GteX (JPMJGX23B2 to A.M.). We thank Xinnian Dong for kindly providing *anac019 anac055 anac072* mutant seeds and Atsushi Takeda for sharing pAT006-Fluc-iG5 and pAT006_Rluc_iFAD2.

## AUTHOR CONTRIBUTIONS

K.T., X.H., and A.M. conceived the idea and supervised the project. W.K., M.N., and K.F. performed most of the experiments. R.H. performed luciferase reporter assays and ChIP-qPCR. W.K. analyzed the data. P.Z. performed stomatal aperture and chlorophyll measurements. Y.D. and K.H. performed phylogenetic analysis. J.Z. measured ABA. Y.T. and D.B. generated transgenic plants. E.N. provided *cyp707a* mutants and advices. R.T.N. and S.M. investigated ABA dynamics using ABA sensors. W.K. and K.T. wrote the manuscript, and all authors contributed to the revision.

---

**Figure S1.**
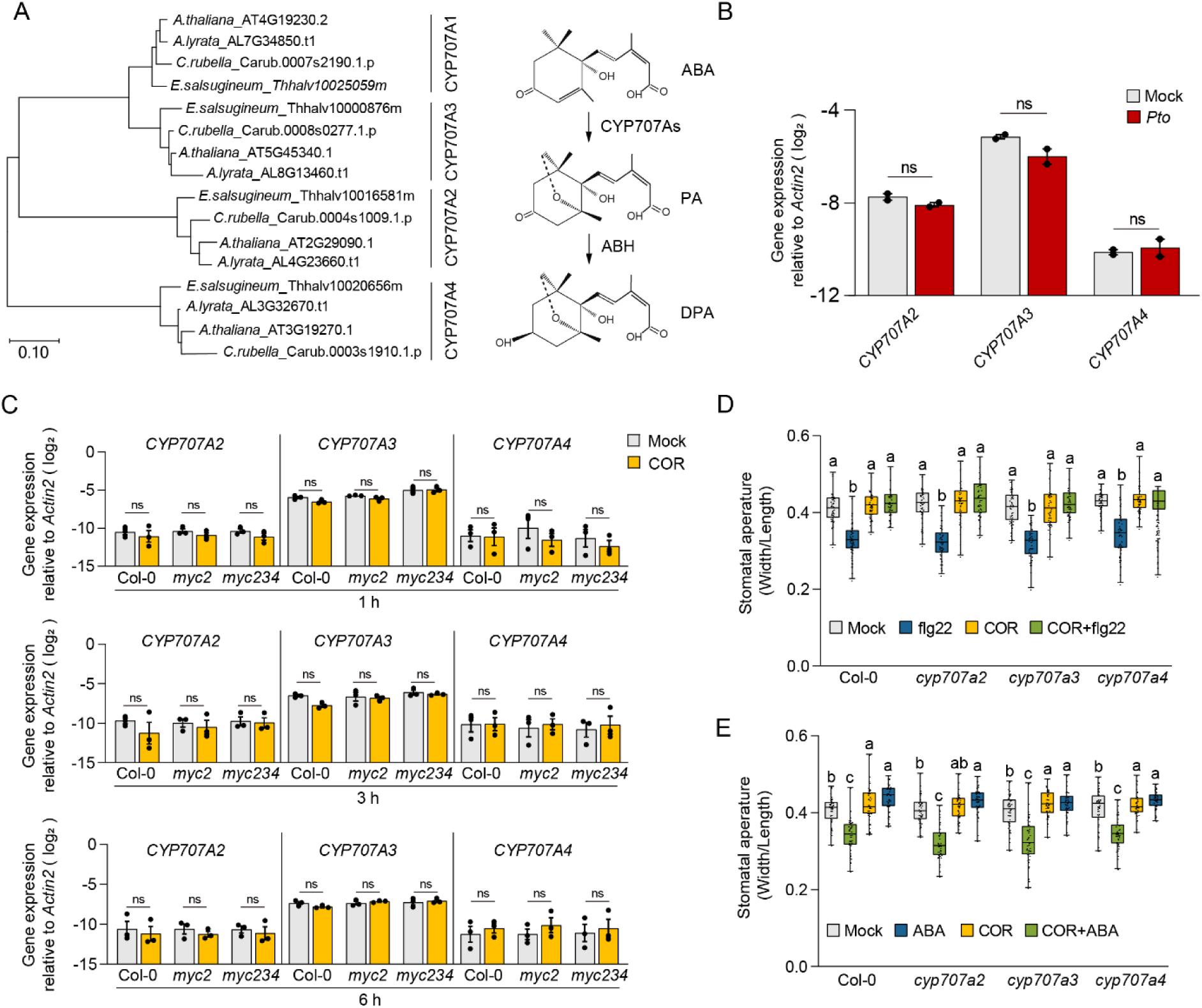
*CYP707A2*, *CYP707A3*, *and CYP707A4* are not essential for COR-mediated inhibition of stomatal closure, related to Figures 1 and 2. (A) The phylogenic trees of CYP707A1, CYP707A2, CYP707A3, and CYP707A4 homologs from the 4 species used in this study. The phylogenic tree was generated using their full-length amino acid sequences and the maximum likelihood method. The scale bar refers to a phylogenetic distance of 0.1 amino acid substitutions per site. ABA catabolism pathway. ABA: abscisic acid. PA: phaseic acid. DPA: dihydrophaseic acid. (B and C) *CYP707As* expression. (B) Leaves of 4 to 5-week-old *A. thaliana* Col-0 WT plants were syringe-infiltrated with mock (water) or *Pto* (OD600=0.001). *CYP707As* expression was measured at 6 hpi by RT-qPCR. (C) Seedlings of 14-day-old *A. thaliana* Col-0 WT and mutant plants were treated with 5 μM COR or mock (0.1% DMSO). *CYP707As* expression was measured at the indicated time points by RT-qPCR. (B and C) Bars represent means and standard errors of log2 expression levels relative to *ACTIN2* calculated from (B) 2 or (C) 3 independent experiments. Asterisks indicate significant differences from mock (\**P <* 0.05, \*\**P <* 0.01, and \*\*\**P* < 0.001, two-tailed Student’s t-tests). (D and E) Stomatal aperture. Stomatal apertures of 4 to 5-week-old *A. thaliana* Col-0 WT and mutant plants were measured after treatment with 5 μM COR or mock (0.1% DMSO) for 30 min, followed by incubation with (D) 5 μM flg22 or mock (water) or (E) 10 μM ABA or mock (0.1% ethanol) for 1 hour. n=48 biological replicates per sample from 2 independent experiments. Results are shown as box plots with boxes displaying the 25th–75th percentiles, the center line indicating the median, and whiskers extending to the minimum and maximum values. All individual data points are overlaid. Statistical analysis was performed within the same genotype, and the results were analyzed using one-way ANOVA followed by Tukey test. Different letters denote a statistically significant difference within a genotype (P < 0.05).

**Figure S2.**
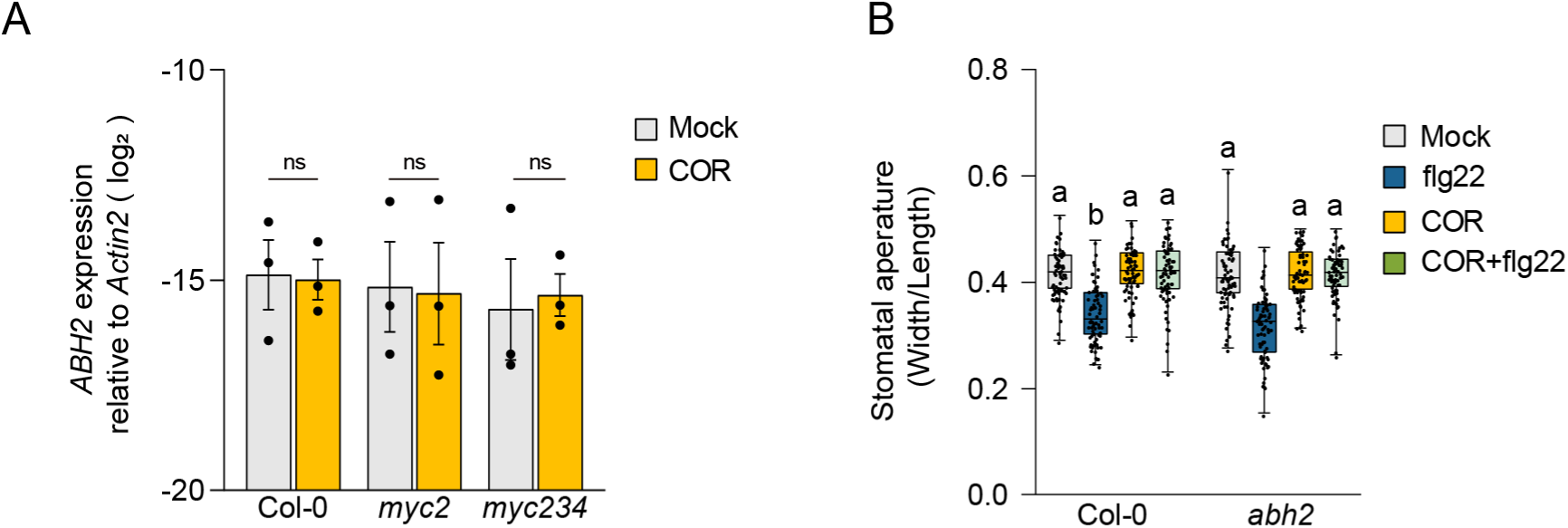
*ABH2* is not required for COR-mediated inhibition of flg22-induced stomatal closure, related to Figure 1. (A) *ABH2* expression. Seedlings of 14-day-old *A. thaliana* Col-0 WT and mutant plants were treated with 5 μM COR or mock (0.1% DMSO). *ABH2* expression was measured at 6 hpi by RT-qPCR. Bars represent means and standard errors of log2 expression levels relative to *ACTIN2* calculated from 3 independent experiments. Asterisks indicate significant differences from mock (\**P <* 0.05, \*\**P <* 0.01, and \*\*\**P* < 0.001, two-tailed Student’s t-tests). (B) Stomatal aperture. Stomatal apertures of 4 to 5-week-old *A. thaliana* Col-0 WT and mutant plants were measured after treatment with 5 μM COR or mock (0.1% DMSO) for 30 min, followed by incubation with 5 μM flg22 or mock (water) for 1 hour. n≥70 biological replicates per sample from 3 independent experiments. Results are shown as box plots with boxes displaying the 25th–75th percentiles, the center line indicating the median, and whiskers extending to the minimum and maximum values. All individual data points are overlaid. Statistical analysis was performed within the same genotype, and the results were analyzed using one-way ANOVA followed by Tukey test. Different letters denote a statistically significant difference within a genotype (P < 0.05).

**Figure S3.**
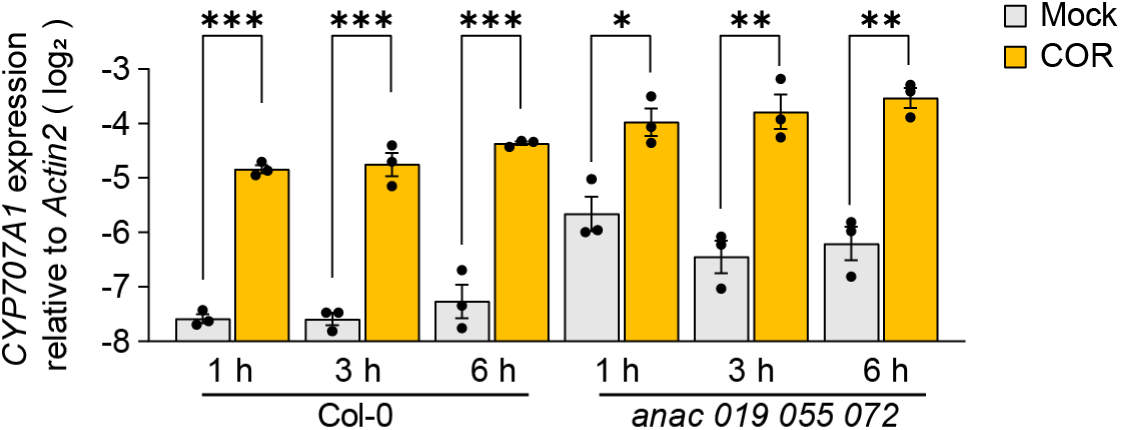
*ANAC019*, *ANAC055*, *and ANAC072* are not required for COR-induced *CYP707A1* expression, related to Figure 2. *CYP707A1* expression. Seedlings of 14-day-old *A. thaliana* Col-0 WT and mutant plants were treated with 5 μM COR or mock (0.1% DMSO). *CYP707A1* expression was measured at the indicated time points by RT-qPCR. (B and C) Bars represent means and standard errors of log2 expression levels relative to *ACTIN2* calculated from 3 independent experiments. Asterisks indicate significant differences from mock (\**P <* 0.05, \*\**P <* 0.01, and \*\*\**P* < 0.001, two-tailed Student’s t-tests).

**Figure S4.**
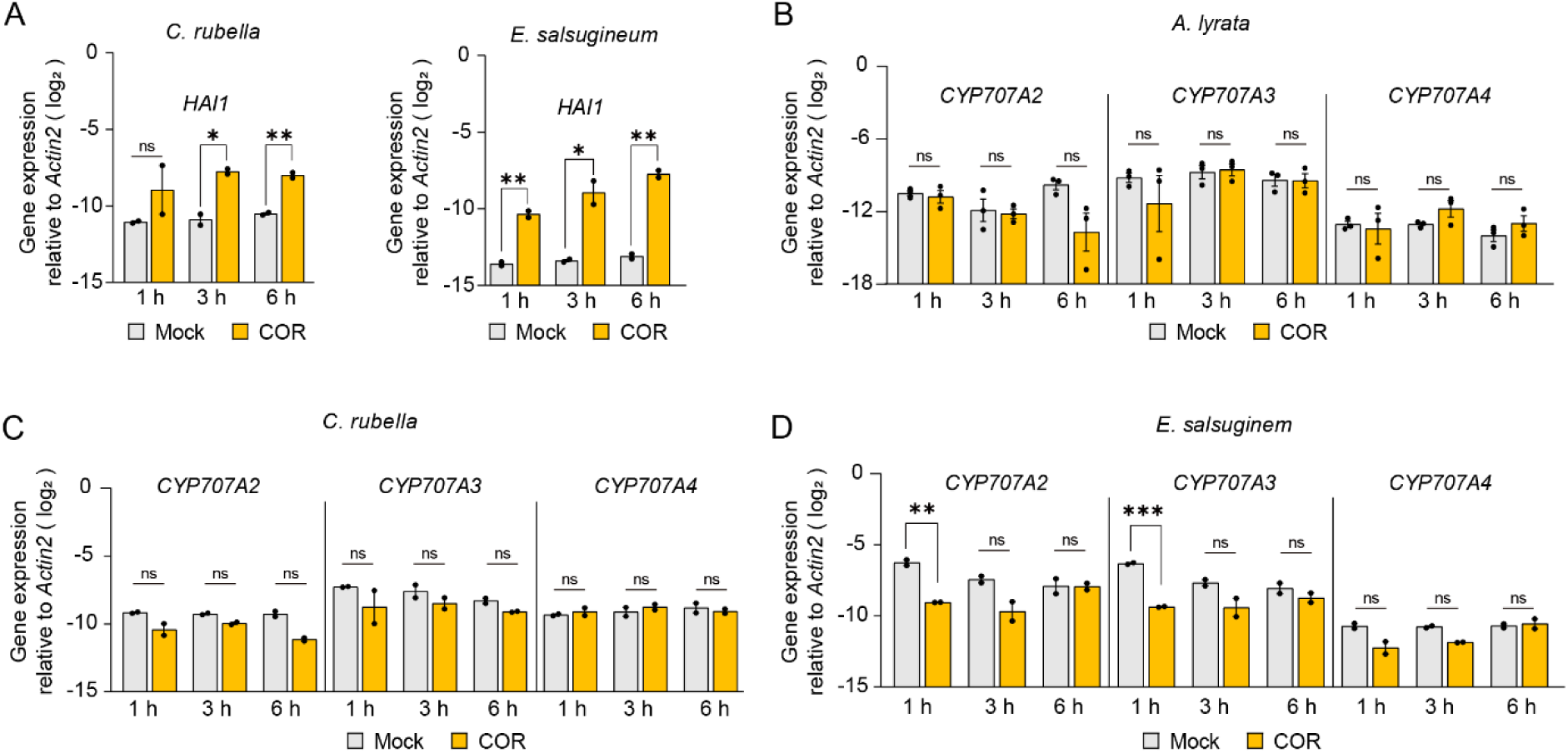
COR does not induce *CYP707A2*, *CYP707A3*, and *CYP707A4* expression in *A. lyrata*, *C. rubella*, and *E. salsugineum*, related to Figure 3. (A to D) Seedlings of 14-day-old *A. lyrata*, *C. rubella*, and *E. salsugineum* plants were treated with 5 μM COR or mock (0.1% DMSO). *HA1*, *CYP707A2*, *CYP707A3*, and *CYP707A4* expression were measured at the indicated time points by RT-qPCR. Bars represent means and standard errors of log2 expression levels relative to *ACTIN2* calculated from (B) 2 or (A, C and D) 3 independent experiments. Asterisks indicate significant differences from mock (\**P <* 0.05, \*\**P <* 0.01, and \*\*\**P* < 0.001, two-tailed Student’s t-tests).

**Figure S5.**
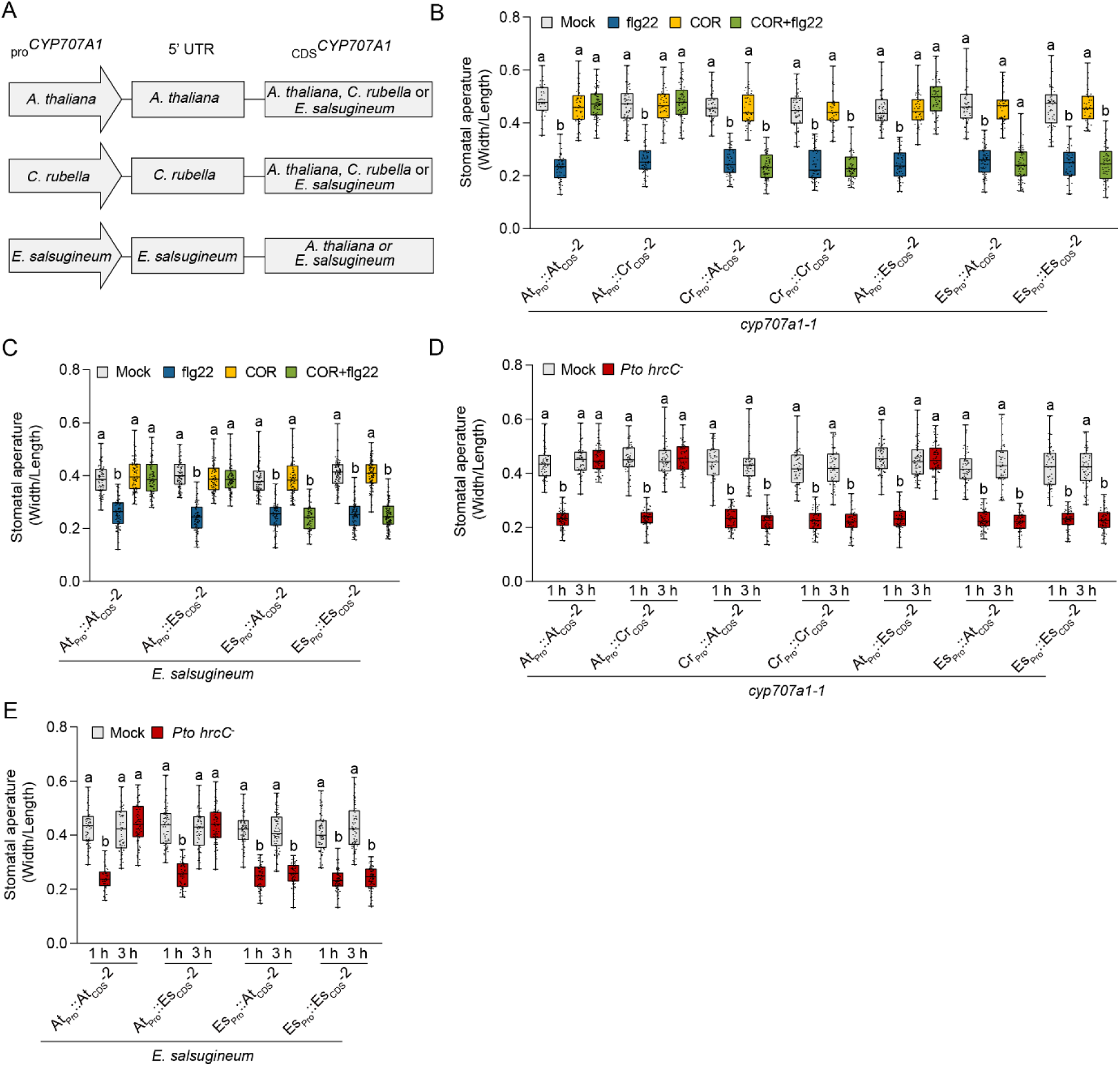
The *CYP707A1* promoter underlies the evolutionary diversification of the COI1-MYC2-CYP707A1 pathway, related to Figure 4. (A) A schematic diagram of the Promoter-Swap transgenic plants. The left column represents the source species of the *CYP707A1* promoter, the middle column represents the source species of the untranslated regions (UTR) associated with the construct, and the right column represents the source species of the *CYP707A1* coding sequence (CDS). (B to E) Stomatal aperture. Stomatal apertures of 4 to 5-week-old *A. thaliana* (B and D) and *E. salsugineum* (C and E) plants were measured after treatment with (B and C) 5 μM COR or mock (0.1% DMSO) for 30 min, followed by incubation with 5 μM flg22 and mock (water); (D and E) at the indicated time points after the treatment with *Pto hrcC^−^* (OD600 = 0.2) or mock (MES buffer). n≥70 biological replicates per sample from 3 independent experiments. Results are shown as box plots with boxes displaying the 25th–75th percentiles, the center line indicating the median, and whiskers extending to the minimum and maximum values. All individual data points are overlaid. Statistical analysis was performed within the same genotype, and the results were analyzed using one-way ANOVA followed by Tukey test. Different letters denote a statistically significant difference within a genotype (P < 0.05).

**Figure S6.**
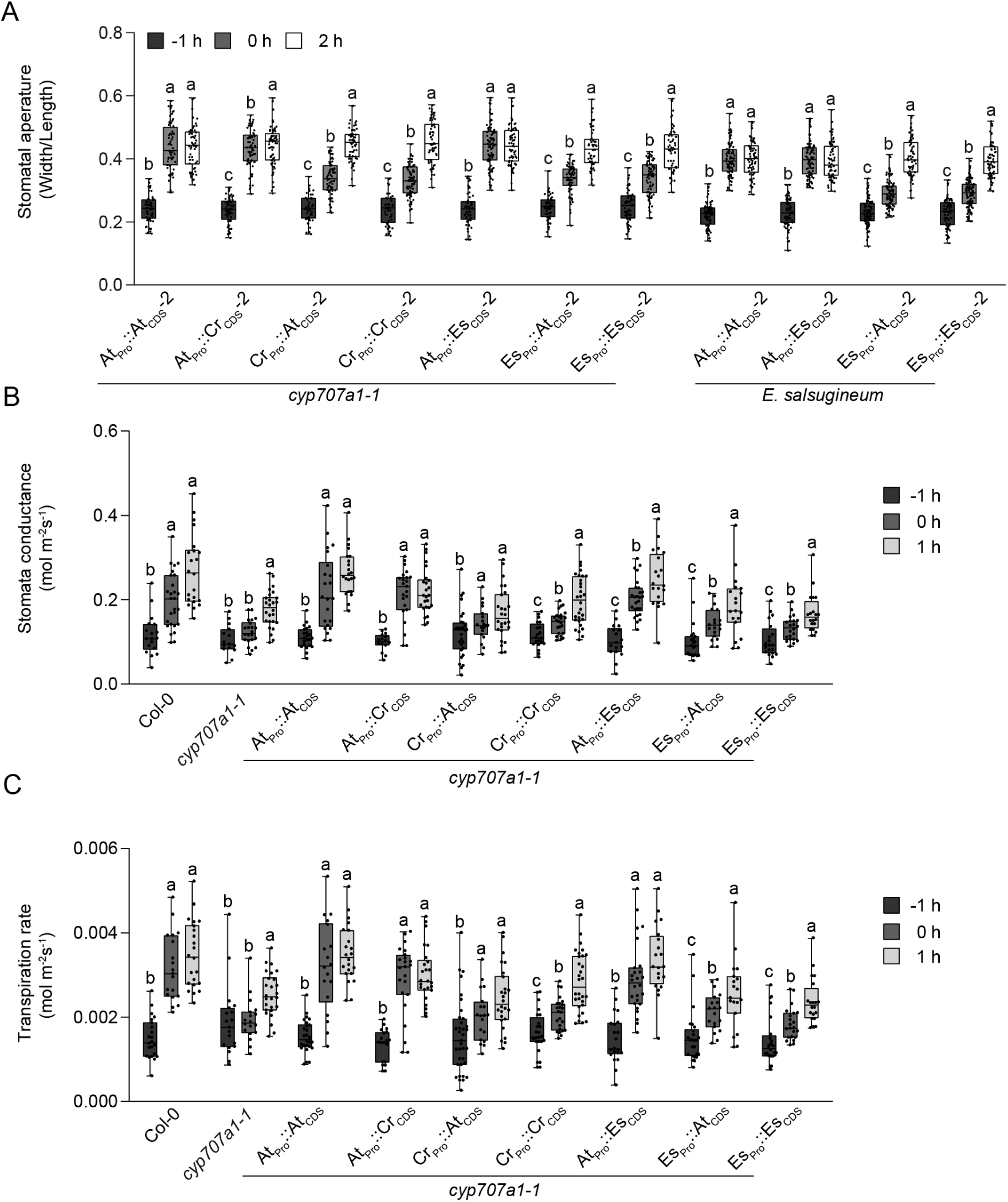
*CYP707A1* regulates light-induced stomata opening at dawn, related to Figure 6. (A) Stomatal aperture. Stomatal apertures of 4 to 5-week-old plants were measured at the indicated time points. n≥70 for each sample from 3 independent experiments. Statistical analysis was performed within the same genotype, and the results were analyzed using one-way ANOVA followed by Tukey test. Different letters denote a statistically significant difference within a genotype (P < 0.05). (B) Stomatal conductance. (C) Transpiration rate. (B and C) Stomata conductance and transpiration rate were determined on leaves of 4-5-week-old plants using a portable photosynthesis system (LI-6800). At least 15 plants per genotype were measured at the designated time from two independent experiments. Statistical analysis was performed within the same genotype, and the results were analyzed using one-way ANOVA followed by Tukey test. Different letters denote a statistically significant difference within a genotype (P < 0.05). (A to D) Results are shown as box plots with boxes displaying the 25th–75th percentiles, the center line indicating the median, and whiskers extending to the minimum and maximum values. All individual data points are overlaid.

**Table S1.**
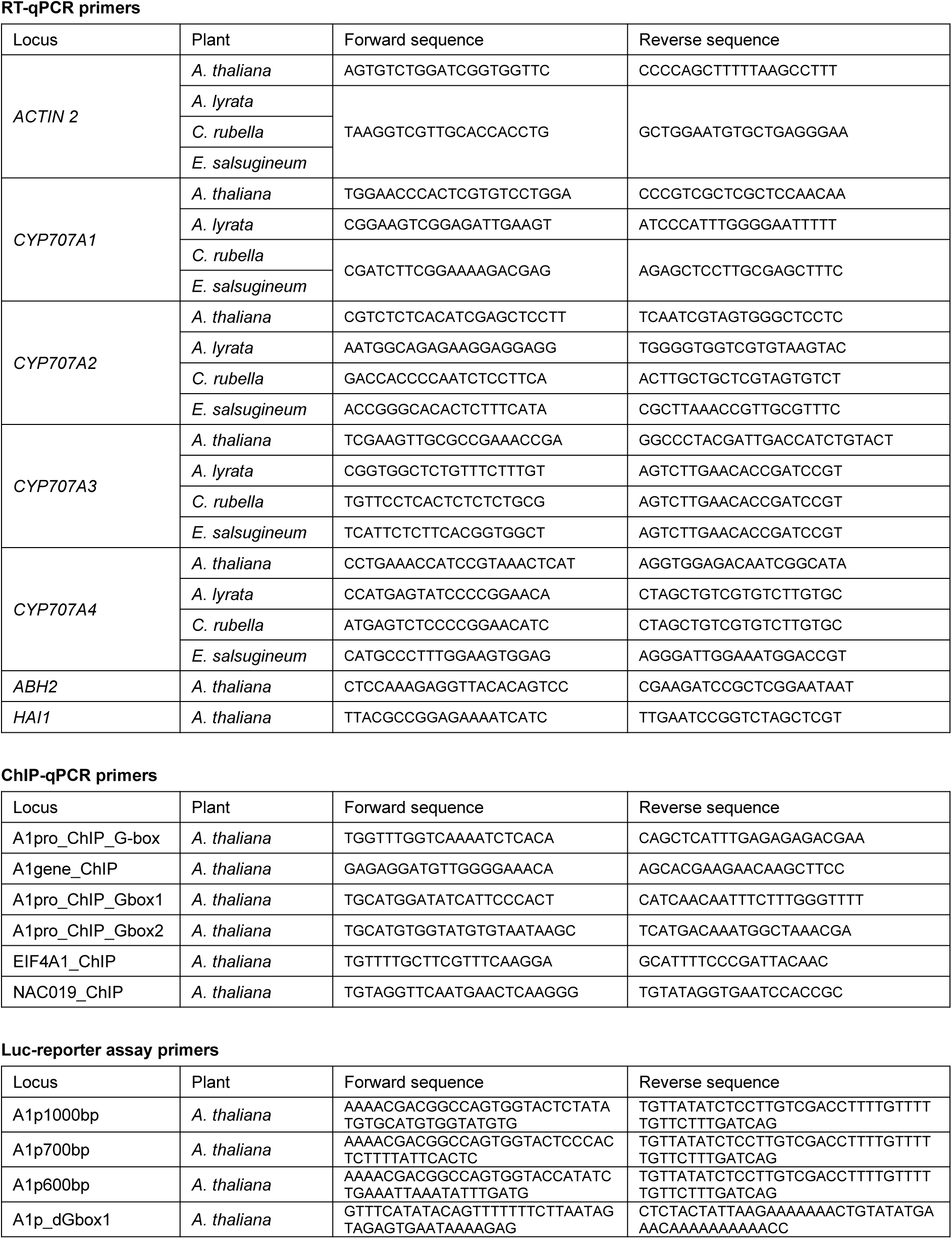
Primers used in this study. Related to STAR Methods.

## STAR ★ METHODS

### KEY RESOURCES TABLE

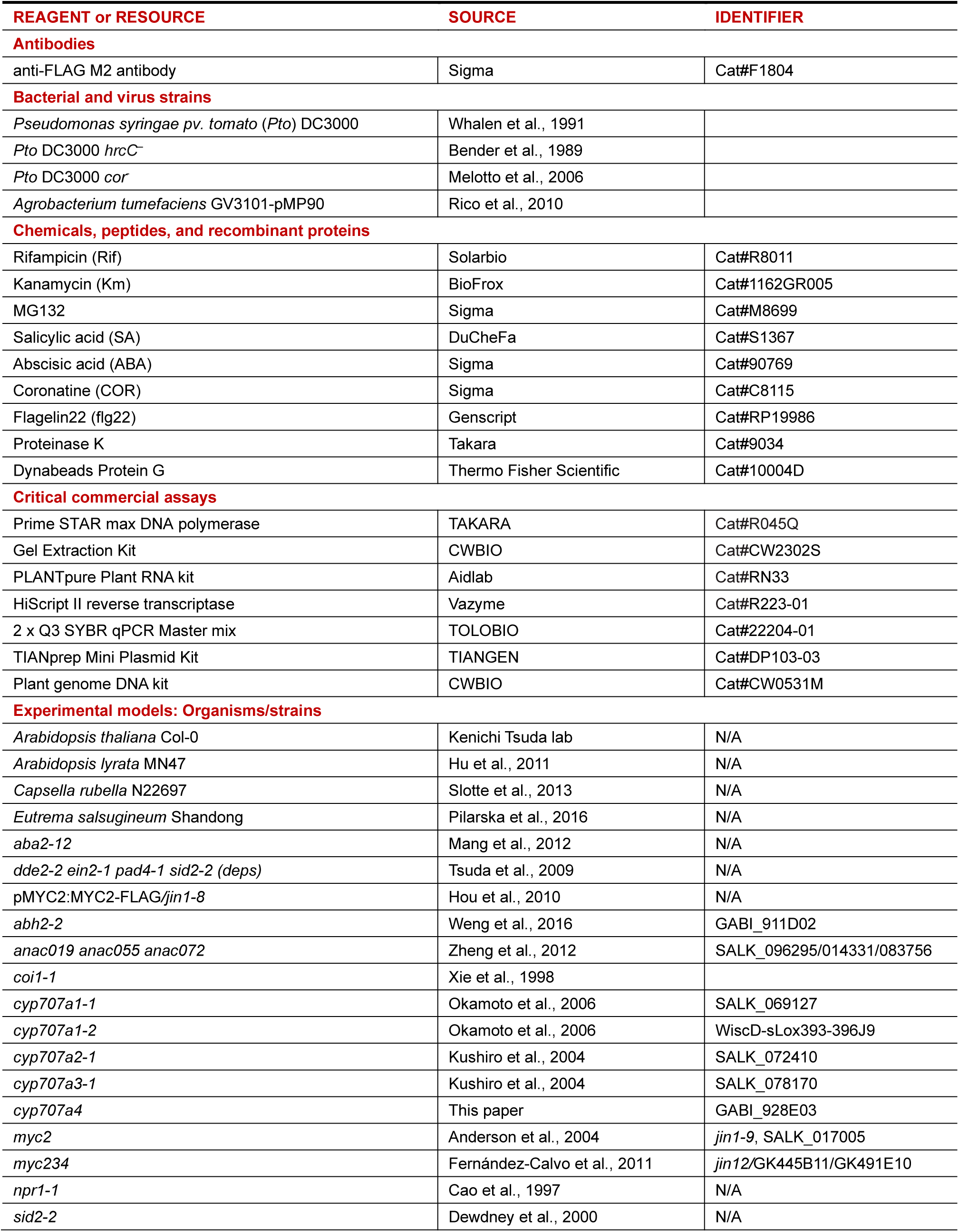
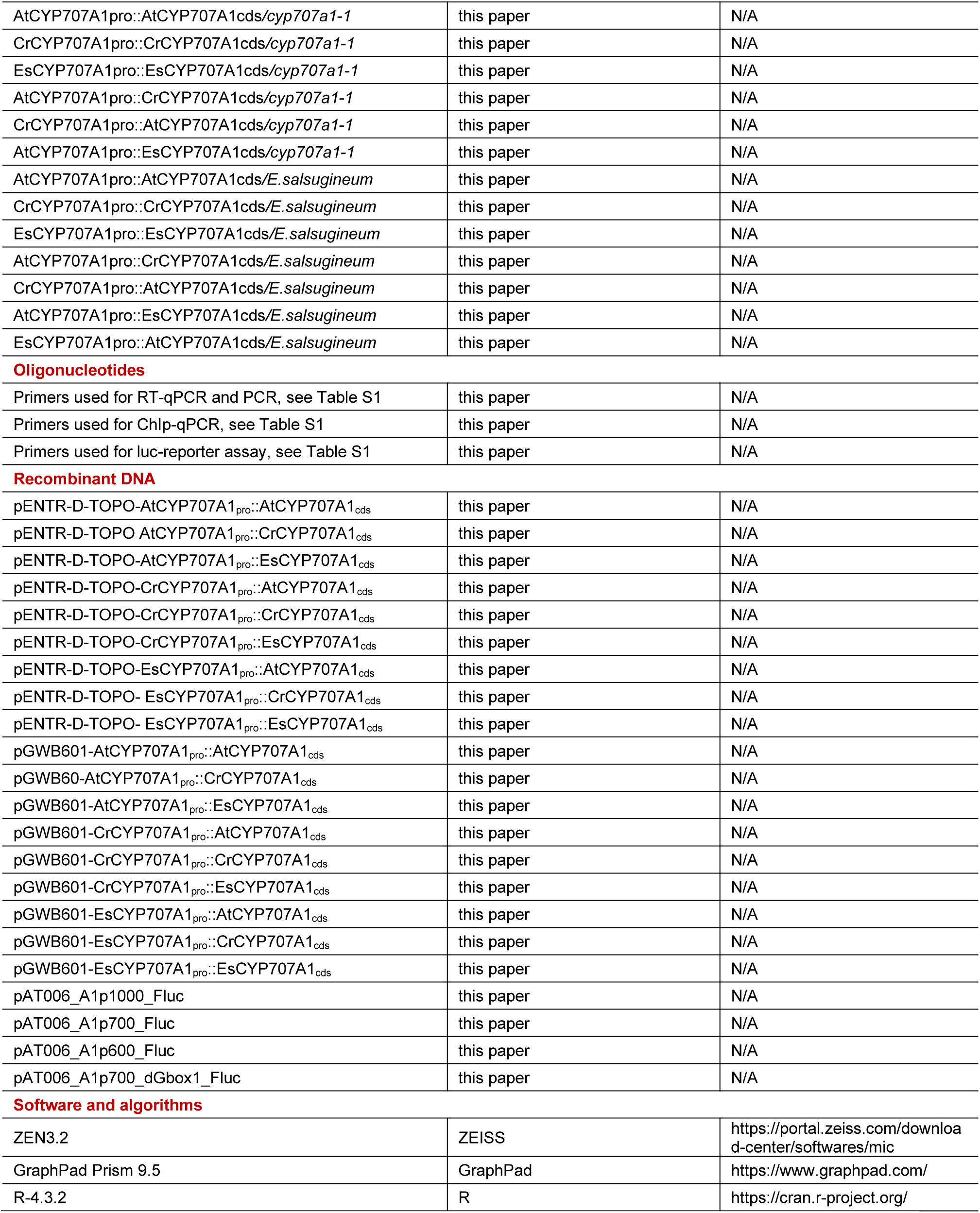

STAR ★ METHODS KEY RESOURCES TABLE
| REAGENT or RESOURCE | SOURCE | IDENTIFIER |
| --- | --- | --- |
| **Antibodies** |  |  |
| anti-FLAG M2 antibody | Sigma | Cat#F1804 |
| **Bacterial and virus strains** |  |  |
| *Pseudomonas syringae* pv. *tomato* (Pto) DC3000 | Whalen et al., 1991 |  |
| Pto DC3000 *hrcC*<sup>−</sup> | Bender et al., 1989 |  |
| Pto DC3000 *cor* | Melotto et al., 2006 |  |
| *Agrobacterium tumefaciens* GV3101-pMP90 | Rico et al., 2010 |  |
| **Chemicals, peptides, and recombinant proteins** |  |  |
| Rifampicin (Rif) | Solarbio | Cat#R8011 |
| Kanamycin (Km) | BioFrox | Cat#1162GR005 |
| MG132 | Sigma | Cat#M8699 |
| Salicylic acid (SA) | DuCheFa | Cat#S1367 |
| Absciscic acid (ABA) | Sigma | Cat#90769 |
| Coronatine (COR) | Sigma | Cat#C8115 |
| Flagelin22 (flg22) | Genscript | Cat#RP19986 |
| Proteinase K | Takara | Cat#9034 |
| Dynabeads Protein G | Thermo Fisher Scientific | Cat#10004D |
| **Critical commercial assays** |  |  |
| Prime STAR max DNA polymerase | TAKARA | Cat#R045Q |
| Gel Extraction Kit | CWBIO | Cat#CW2302S |
| PLANTpure Plant RNA kit | Aidlab | Cat#RN33 |
| HiScript II reverse transcriptase | Vazyme | Cat#R223-01 |
| 2 x Q3 SYBR qPCR Master mix | TOLOBIO | Cat#22204-01 |
| TIANprep Mini Plasmid Kit | TIANGEN | Cat#DP103-03 |
| Plant genome DNA kit | CWBIO | Cat#CW0531M |
| **Experimental models: Organisms/strains** |  |  |
| *Arabidopsis thaliana* Col-0 | Kenichi Tsuda lab | N/A |
| *Arabidopsis lyrata* MN47 | Hu et al., 2011 | N/A |
| *Capsella rubella* N22697 | Slotte et al., 2013 | N/A |
| *Eutrema salsugineum* Shandong | Pilarska et al., 2016 | N/A |
| *aba2-12* | Mang et al., 2012 | N/A |
| *dde2-2 ein2-1 pad4-1 sid2-2 (deps)* | Tsuda et al., 2009 | N/A |
| pMYC2:MYC2-FLAG/*jin1-8* | Hou et al., 2010 | N/A |
| *abh2-2* | Weng et al., 2016 | GABI_911D02 |
| *anac019 anac055 anac072* | Zheng et al., 2012 | SALK_096295/014331/083756 |
| *coi1-1* | Xie et al., 1998 |  |
| *cyp707a1-1* | Okamoto et al., 2006 | SALK_069127 |
| *cyp707a1-2* | Okamoto et al., 2006 | WiscD-sLox393-396J9 |
| *cyp707a2-1* | Kushiro et al., 2004 | SALK_072410 |
| *cyp707a3-1* | Kushiro et al., 2004 | SALK_078170 |
| *cyp707a4* | This paper | GABI_928E03 |
| *myc2* | Anderson et al., 2004 | *jin1-9*, SALK_017005 |
| *myc234* | Fernández-Calvo et al., 2011 | *jin12/GK445B11/GK491E10* |
| *npr1-1* | Cao et al., 1997 | N/A |
| *sid2-2* | Dewdney et al., 2000 | N/A |
| AtCYP707A1<sub>pro</sub>::AtCYP707A1<sub>cds</sub>/*cyp707a1-1* | this paper | N/A |
| CrCYP707A1<sub>pro</sub>::CrCYP707A1<sub>cds</sub>/*cyp707a1-1* | this paper | N/A |
| EsCYP707A1<sub>pro</sub>::EsCYP707A1<sub>cds</sub>/*cyp707a1-1* | this paper | N/A |
| AtCYP707A1<sub>pro</sub>::CrCYP707A1<sub>cds</sub>/*cyp707a1-1* | this paper | N/A |
| CrCYP707A1<sub>pro</sub>::AtCYP707A1<sub>cds</sub>/*cyp707a1-1* | this paper | N/A |
| AtCYP707A1<sub>pro</sub>::EsCYP707A1<sub>cds</sub>/*cyp707a1-1* | this paper | N/A |
| AtCYP707A1<sub>pro</sub>::AtCYP707A1<sub>cds</sub>/*E.salsugineum* | this paper | N/A |
| CrCYP707A1<sub>pro</sub>::CrCYP707A1<sub>cds</sub>/*E.salsugineum* | this paper | N/A |
| EsCYP707A1<sub>pro</sub>::EsCYP707A1<sub>cds</sub>/*E.salsugineum* | this paper | N/A |
| AtCYP707A1<sub>pro</sub>::CrCYP707A1<sub>cds</sub>/*E.salsugineum* | this paper | N/A |
| CrCYP707A1<sub>pro</sub>::AtCYP707A1<sub>cds</sub>/*E.salsugineum* | this paper | N/A |
| AtCYP707A1<sub>pro</sub>::EsCYP707A1<sub>cds</sub>/*E.salsugineum* | this paper | N/A |
| EsCYP707A1<sub>pro</sub>::AtCYP707A1<sub>cds</sub>/*E.salsugineum* | this paper | N/A |
| **Oligonucleotides** |  |  |
| Primers used for RT-qPCR and PCR, see Table S1 | this paper | N/A |
| Primers used for ChIP-qPCR, see Table S1 | this paper | N/A |
| Primers used for luc-reporter assay, see Table S1 | this paper | N/A |
| **Recombinant DNA** |  |  |
| pENTR-D-TOPO-AtCYP707A1<sub>pro</sub>::AtCYP707A1<sub>cds</sub> | this paper | N/A |
| pENTR-D-TOPO AtCYP707A1<sub>pro</sub>::CrCYP707A1<sub>cds</sub> | this paper | N/A |
| pENTR-D-TOPO-AtCYP707A1<sub>pro</sub>::EsCYP707A1<sub>cds</sub> | this paper | N/A |
| pENTR-D-TOPO-CrCYP707A1<sub>pro</sub>::AtCYP707A1<sub>cds</sub> | this paper | N/A |
| pENTR-D-TOPO-CrCYP707A1<sub>pro</sub>::CrCYP707A1<sub>cds</sub> | this paper | N/A |
| pENTR-D-TOPO-CrCYP707A1<sub>pro</sub>::EsCYP707A1<sub>cds</sub> | this paper | N/A |
| pENTR-D-TOPO-EsCYP707A1<sub>pro</sub>::AtCYP707A1<sub>cds</sub> | this paper | N/A |
| pENTR-D-TOPO- EsCYP707A1<sub>pro</sub>::CrCYP707A1<sub>cds</sub> | this paper | N/A |
| pENTR-D-TOPO- EsCYP707A1<sub>pro</sub>::EsCYP707A1<sub>cds</sub> | this paper | N/A |
| pGWB601-AtCYP707A1<sub>pro</sub>::AtCYP707A1<sub>cds</sub> | this paper | N/A |
| pGWB601-AtCYP707A1<sub>pro</sub>::CrCYP707A1<sub>cds</sub> | this paper | N/A |
| pGWB601-AtCYP707A1<sub>pro</sub>::EsCYP707A1<sub>cds</sub> | this paper | N/A |
| pGWB601-CrCYP707A1<sub>pro</sub>::AtCYP707A1<sub>cds</sub> | this paper | N/A |
| pGWB601-CrCYP707A1<sub>pro</sub>::CrCYP707A1<sub>cds</sub> | this paper | N/A |
| pGWB601-CrCYP707A1<sub>pro</sub>::EsCYP707A1<sub>cds</sub> | this paper | N/A |
| pGWB601-EsCYP707A1<sub>pro</sub>::AtCYP707A1<sub>cds</sub> | this paper | N/A |
| pGWB601-EsCYP707A1<sub>pro</sub>::CrCYP707A1<sub>cds</sub> | this paper | N/A |
| pGWB601-EsCYP707A1<sub>pro</sub>::EsCYP707A1<sub>cds</sub> | this paper | N/A |
| pAT006_A1p1000_Fluc | this paper | N/A |
| pAT006_A1p700_Fluc | this paper | N/A |
| pAT006_A1p600_Fluc | this paper | N/A |
| pAT006_A1p700_dGbox1_Fluc | this paper | N/A |
| **Software and algorithms** |  |  |
| ZEN3.2 | ZEISS | https://portal.zeiss.com/download-center/software/mic |
| GraphPad Prism 9.5 | GraphPad | https://www.graphpad.com/ |
| R-4.3.2 | R | https://cran.r-project.org/ |

### RESOURCE AVAILABILITY

#### Lead contact

Further information and requests for resources and reagents should be directed to and will be fulfilled by the lead contact, Kenichi Tsuda

#### Materials availability

Unique reagents generated in this study are available from the lead contact with a completed Materials Transfer Agreement.

### EXPERIMENTAL MODEL AND STUDY PARTICIPANT DETAILS

All cell lines and source materials used in this study are listed in the key resources table.

#### METHOD DETAILS

##### Plant materials and growth conditions

*A. lyrate* accession MN47, *C. rubella* N22697, *E. salsugineum* Shandong, *A. thaliana* accession Col-0, mutants, and transgenic plants used in this study were grown in soil for four to five weeks in growth chambers (22°C, 60% relative humidity, 200 μE m-2s-1 light with a 12-hr light/12-hr dark) for stomatal aperture, stomatal conductance, and pathogen assays. For COR-induced gene expression analysis, seeds were sterilized in a bleach solution for three minutes, followed by five washes with sterile water. After two days of stratification at 4°C in the dark, seeds were sown on plates containing half-strength Murashige and Skoog medium (½MS) supplemented with 1% sucrose and 0.5% agar (pH 5.7) and grown for 13 days in the same growth chamber. Seedlings were transferred to liquid ½ MS-medium one day before experiments. The *aba2-12*, *dde2-2 ein2-1 pad4-1 sid2-2* (*deps*), pMYC2::MYC2-FLAG*/jin1-8*, *abh2-12* (GABI_911D02), *anac019 anac055 anac072* (SALK_096295/014331/083756), *coi1-1*, *cyp707a1-1* (SALK_069127), *cyp707a1-2* (WiscD-sLox393-396J9), *cyp707a2-1*, *cyp707a3-1*, *myc2* (*jin1-9*, SALK_017005), *myc234 (jin1-2/*GK445B11/GK491E10), *npr1-1*, *sid2-2* were previously described. Transgenic plants were generated using floral dip transformation63 and selected using BASTA spray. Homozygous T3 transgenic plants were used for experiments.

##### Bacterial cultivation

Details of the bacterial strains used are provided in the key resources table. *Pto*, *Pto hrcC–*, and *Pto cor–* strains were cultured on KB agar plates with 40 μg/ml rifampicin at 28°C for two days. Colonies were resuspended in 900 μl of KB liquid medium, followed by inoculation into 8 ml KB liquid medium with 40 μg/ml rifampicin and incubation at 28°C for 12-15 hours. The bacterial suspension was harvested by centrifugation and washed with sterile water.

##### Bacterial growth assay

Leaves of four to five-week-old plants were infiltrated with a bacterial suspension (OD600 = 0.0002) or sprayed (OD600 = 0.2) using Silwet L-77 (0.04%) as a surfactant. Leaf discs were collected after two days (infiltration) or three days (spraying), homogenized with sterile 10 mM MgCl2, serially diluted, and plated on KB agar supplemented with 40 μg/ml rifampicin for colony counting.

##### Bacterial invasion assay

Plants were kept for at least three hours under light before infection. Fully-expanded leaves were cut and placed on the abaxial side onto the bacterial suspension. Afterward, the leaves were washed with 5% hydrogen peroxide, and homogenized for colony counting. Log10-transformed colony-forming units (CFU) per cm2 leaf surface area were calculated.

##### RT-qPCR

Ten seedlings were transferred to 1/2 MS liquid medium and treated with DMSO (0.1%) or 5 μM COR. Approximately five leaves were harvested for light-induced stomatal opening experiments. For gene expression response to bacteria, leaves of four to five-week-old plants were syringe-infiltrated with mock (water) or bacteria at OD600=0.001, and harvested at 6 hours post infiltration. Total RNA was isolated with a PLANTpure Plant RNA kit. RNA concentration was measured and adjusted to 100 ng/μL with nuclease-free water for complementary DNA (cDNA) synthesis. HiScript II reverse transcriptase was used for cDNA synthesis. RT-qPCR reaction was performed with 2 x Q3 SYBR qPCR Master Mix by using an IQ5 real-time PCR Thermocycler (Bio-Rad). Fragments of target genes were amplified using the primers listed in Table S1, and specific gene expression was normalized to *ACTIN2*.

##### ChIP-qPCR analysis

Chromatin immunoprecipitation was carried out essentially according to Nomoto et al. (2021).64 Briefly, two-week-old *A. thaliana* seedlings grown in liquid half-strength MS medium supplemented with 1% sucrose were treated with 5 μM COR for three hours. Mock-treated seedlings were also harvested. After fixation, tissues were frozen in liquid nitrogen and stored at -80°C. Frozen tissues (∼1 g) were ground in liquid nitrogen and suspended in a 2.5 mL nuclei extraction buffer [10 mM Tris-HCl, pH 8.0, 0.25 M sucrose, 10 mM MgCl2, 40 mM β-mercaptoethanol, and complete protease inhibitor cocktail (Roche)]. The suspension was filtered through two layers of Miracloth and centrifuged at 17,500×g at 4°C for 5 min. The pellet was resuspended in 75 μL of nuclei lysis buffer [50 mM Tris-HCl, pH 8.0, 10 mM EDTA, 1% (w/v) SDS]. After incubation at room temperature for 20 min and then on ice for 10 min, 225 μL of ChIP dilution buffer [50 mM Tris-HCl, pH 8.0, 10 mM EDTA, 1% (w/v) SDS] was added. The sample was then transferred to a new 1.5 mL tube and sonicated at high power for 10 cycles of 30 s ON/30 s OFF using BIORUPTOR II to produce DNA fragments. After addition of 375 μL of ChIP dilution buffer, 200 μL of ChIP dilution buffer with Triton, and 35 μL of 20% (w/v) Triton X-100, the sample was centrifuged at 17,500×g at 4°C for 5 min. The supernatant was used as the starting material for chromatin immunoprecipitation using 5 μg of anti-FLAG M2 antibody and 75 μL of Dynabeads Protein G for 2 h at 4°C with gentle rotation. In contrast, 18 μL of the supernatant was kept as the input sample. For the input controls, 82 μL of elution buffer was added. Both the ChIP and input samples were mixed with 6 μl of 5 M NaCl and incubated at 65°C overnight, followed by digestion with Proteinase K at 37°C for 1 h. DNA was purified using NucleoSpin Gel and PCR cleanup kit with Buffer NTB (MACHEREY-NAGEL). Quantitative PCR was performed using primers listed in Table S1. The percentage of input values of the ChIP DNA was further normalized over the value obtained for the *eIF4A* promoter (At3g13920).

##### Luciferase reporter assay

The 1000 bp, 700 bp, and 600 bp sequences upstream of the start codon of the *CYP707A1* gene were amplified by PCR from *A. thaliana* genomic DNA. These PCR products were assembled by NEBuilder HiFi DNA assembly reactions (NEB) into pAT006-Fluc-iG5 that had been digested with KpnI and SalI, creating pAT006_A1p1000_Fluc, pAT006_A1p700_Fluc, and pAT006_A1p600_Fluc, respectively. To construct pAT006_A1p700_dGbox1_Fluc, two PCR fragments were amplified from pAT006_A1p1000_Fluc using primer pairs A1p1000bp_F plus A1p_dGbox1_R and A1p_dGbox1_F plus A1p1000bp_R. These two PCR fragments were then used as templates for PCR using A1p700bp_F and A1p1000bp_R, followed by a NEBuilder HiFi DNA assembly reaction with KpnI/SalI-digested pAT006-Fluc-iG5. Primers used for the construction of luciferase reporters are listed in Table S1. *CYP707A1* promoter activity assay was performed using *Agrobacterium*-mediated transient expression in *A. thaliana dde2 ein2 pad4 sid2* (*deps*) mutant.38 Firefly luciferase (Fluc) reporter constructs and the internal control construct pAT006_Rluc_iFAD2 carrying *Renilla* luciferase (Rluc) gene under the control of a cauliflower mosaic virus 35S promoter were introduced into *A. tumefaciens* strain GV3101 (pMP90). *Agrobacterium* strains carrying Fluc reporter constructs or pAT006_Rluc_iFAD2 were mixed at a 10:1 ratio at OD600 of 0.1 and infiltrated into leaves of 3 to 4-week-old *deps* plants. Two days later, the infiltrated leaves were treated with 5 μM coronatine and harvested 3 hours later. Fluc and Rluc activities were measured using the Dual-Luciferase reporter assay system (Promega) on the GloMax Navigator Microplate Luminometer (Promega).

##### Stomatal aperture measurement and analysis

Four to five-week-old plants were kept under light (200 μE m-2s-1) for at least 3 h to ensure stomata were open before starting experiments. Similarly developed leaf epidermis was peeled and immersed in MES buffer with 5 μM COR or mock (0.1% DMSO) for 30 min, followed by incubation with 5 μM flg22 or mock (water); 10 μM ABA or mock (0.1% ethanol); 100 μl SA or mock (water) for 1 h. For measuring stomatal response to bacteria, the peeled leaf epidermis was immersed in bacterial suspension (OD600 = 0.2 in MES buffer) for 1 h and 3 h. Pictures of the leaf epidermis were taken by OLYMPUS IX71SP8 (OLYMPUS, Japan) under normal light. For measuring stomatal aperture in the morning, pictures were taken 1-0.5 h before and 0.5-1 h, 1.5-2 h, and 2.5-3 h after light on. The stomatal aperture was measured by taking ratios of width and length using ZEN software.

##### Stomatal conductance and transpiration rate analysis

Stomatal conductance and transpiration rate were measured in the leaves of four to five-week-old plants using a portable photosynthesis system (LI-6800). The measurement conditions were 200 μmol m−2 s−1 light intensity, 55-60% relative humidity, 22°C, and 400 ppm CO2. Measurements were recorded 1-0.5 h before and 0.5-1 h, 1.5-2 h after light exposure. Data were recorded until the transpiration rate and stomatal conductance curves reach a steady state.

##### Stomatal density analysis

To measure stomatal density, leaves of 4 to 5-week-old plants were selected. Print the abaxial surfaces on clear tape, and remove green tissue with tweezers to expose the stomatal imprints. And then the stomatal imprints were mounted on the glass slide. The number of stomata was counted using the images captured at 40× magnification, in a field of 0236 mm−2 using a fluorescence microscope (OLYMPUS IX71SP8). The stomatal density (SD) was calculated using the formula: stomatal density = the number of stomata/area in view.

##### Chlorophyll content detection

Plants grown under normal light conditions (200 μE m-2s-1 light with a 12-hr light/12-hr dark) for three weeks were transferred to a high-intensity environment (600 μE m-2s-1 light with a 12-hr light/12-hr dark). After two days of treatment, chlorophyll content was measured using a SPAD meter. For each plant, three similarly developed leaves were selected for SPAD measurements, and the average of these three values was calculated to represent the chlorophyll content of the plant.

##### Molecular cloning

The *CYP707A1* promoter was amplified from *Arabidopsis thaliana*, *Arabidopsis lyrate*, *Capsella rubella*, and *Eutrema salsugineum* genomic DNA respectively. The coding sequence of the *CYP707A1* gene was amplified from the cDNA of the plants mentioned above. The size of PCR products was checked by electrophoresis and the gel bands were cut and purified by Gel Extraction Kit (CWBIO, China). To generate the pENTR-D-TOPO donor vector, containing the *CYP707A1* gene from various plant accessions driven by the *AtCYP707A1* promoter, *CrCYP707A1* promoter, and *EsCYP707A1* promoter, the promoter (1 kb upstream of the start codon) and code sequencing of each gene was amplified, and ligated into pENTR-D-TOPO to obtain pENTR-D-TOPO-AtCYP707A1pro::AtCYP707A1cds, pENTR-D-TOPO AtCYP707A1pro::CrCYP707A1cds, pENTR-D-TOPO-AtCYP707A1pro::EsCYP707A1cds, pENTR-D-TOPO-CrCYP707A1pro::AtCYP707A1cds, pENTR-D-TOPO-CrCYP707A1pro::CrCYP707A1cds, pENTR-D-TOPO-CrCYP707A1pro::EsCYP707A1cds, pENTR-D-TOPO-EsCYP707A1pro::AtCYP707A1cds, pENTR-D-TOPO-EsCYP707A1pro::CrCYP707A1cds, pENTR-D-TOPO- EsCYP707A1pro::EsCYP707A1cds. Ligation of PCR products and entry vector pENTR-D-TOPO was performed with In-Fusion HD Cloning Plus, using the reaction mix (5X In-fusion HD Enzyme Premix, 2 μl; 50 ng/μl Linearized vector, 2 μl; 50 ng/μl purified DNA fragment 1, 2 μl; 50 ng/μl purified DNA fragment 2, 2 μl; Sterile H2O, 2 μl). The ligation mixture was incubated for 15 min at 50°C. Afterward, competent *E. coli* cells were transformed with the plasmids that were mentioned above, and positively transformed colonies were picked on LB medium plate with appropriate antibiotics. Isolated plasmids were analyzed by Sanger sequencing to check whether they were the correct plasmid. The destination vector was made by LR reaction from donor vector (pENTR-D-TOPO ligated with fragments) to acceptor vector pGWB601 containing BASTA resistance. The LR reaction mixture was incubated up to 18 h at 25°C. After incubation, 1 μl of proteinase K solution was added, further incubated for 10 min at 37°C, and then transformed into Stellar competent *E. coli* cells DH5α. The sequences of all genes or promoters were verified by Sanger sequencing. The destination plasmids were transformed into the *Agrobacterium tumefaciens* strain and then introduced into plants using floral dip transformation.

##### Phylogenetic tree construction

The construction of the gene phylogenetic tree begins with retrieving the amino acid sequences of the *CYP707A* family from *A. thaliana* using the Phytozome database. Subsequently, homologous *CYP707A* sequences from additional species were identified through BLAST searches. The collected sequences are imported into MEGA software, where the maximum likelihood method is employed to construct the phylogenetic tree. A text file listing the species names is first prepared to construct the species phylogenetic tree. This file is uploaded to the TimeTree platform, which utilizes established evolutionary data to generate a species phylogenetic tree, illustrating the evolutionary relationships and divergence times among the species.

##### Media and reagents

1/2 Murashige & Skoog (MS) (pH 5.7): 2.2 g/l Murashige and Skoog medium include vitamins, 1 % (w/v) sucrose, 0.5 % (w/v) plant agar. Sterilizing bleach solution: 1.5% (v/v) NaClO, 0.02% (v/v) TritonX-100. King’s B (KB) (pH 6.9): 10 g/l Bacto Proteose peptone No. 3, 15 g/l Glycerol, 1.5 g/l K2HPO4, 15 g/l Bacto agar. Luria-Bertani (LB) (pH 7.0): 1% (w/v) Trypton, 1 % (w/v) NaCl, 0.5 % (w/v) yeast extract, 1.5 % (w/v) agar. MES buffer (pH 6.5) 25 mM MES-KOH, 10 mM KCl. Lysis buffer (pH 8.0): 50 mM Tris-HCl, 2 mM EDTA, 150 mM NaCl, 1% Triton X-100, 50 μM MG132 (Sigma).

##### Statistical analyses

Statistical significances were analyzed as written in each figure legend using the R and GraphPad Prism 9.5 software.

